# Sex differences distinguish performance in four object recognition-based memory tasks in the Pink1-/- rat model of Parkinson’s disease

**DOI:** 10.1101/2025.02.19.639089

**Authors:** Aveena M. Desai, Oluwagbohunmi A. Aje, Mary F. Kritzer

**Affiliations:** Department of Neurobiology and Behavior, Stony Brook University, Stony Brook New York 11794-5230, USA

**Keywords:** Non-motor deficits, PTEN-induced putative kinase1, Episodic memory, Spatial cognition, Novel object recognition, Novel object location, Object-in-Place

## Abstract

Many patients with Parkinson’s disease (PD) experience early, sometimes prodromal non-motor deficits involving cognition and memory. These so-called mild cognitive impairments hold dire predictions for future risk of freezing, falls and developing PD-related dementia. Moreover, due to a dearth of effective treatments, these symptoms persist and progressively worsen. Thus, there is an urgent need to better understand and better treat these debilitating signs. Sex differences in incidence, severity and treatment sensitivities predict that the answers to these questions are sex-specific. The work presented here highlights new ways in which rats with knockout of PTEN-induced putative kinase 1 gene (Pink1-/-) emulate PD’s mild cognitive deficits and their clinical sex differences. Specifically, longitudinal behavioral testing confirmed that male Pink1-/- rats developed significant deficits in Novel Object Recognition and Novel Object Location tasks by 5 months old but that female Pink1-/- were unimpaired in these and the Object-in-Place task through 12 months of age. Further, What, Where, When Episodic-like Memory testing identified enduring deficits in all three memory domains in Pink1-/- males by 3 months of age whereas in Pink1-/- females, non-significant impairments emerged at 7 months of age and progressed to significant memory deficits by 12 months of age. Together, these data show that Pink1-/- rats model the generally greater vulnerability of male PD patients to cognitive and memory deficits in PD, the growing risk for higher order deficits in female patients as they age, and features including early onset that distinguish episodic memory impairments from other at-risk processes in this disorder.

## 1.0 INTRODUCTION

Parkinson’s Disease (PD) is a neurodegenerative disorder characterized by diverse motor symptoms, including bradykinesia and postural instability(*1–3*) and by non-motor deficits, including disturbances of sleep, sensory perceptions, affect, and autonomic function(*4–6*). In addition, at the time of diagnosis roughly 20% of patients also experience cognitive impairments, typically in domains of executive function, attention, visuospatial cognition and/or episodic or other types of memory (*7–12*). The clinical term “mild cognitive deficits” is used to distinguish these early appearing symptoms from later presenting disturbances associated with PD-related dementia. However, these so-named mild deficits are described by patients and caregivers as distressing and disabling and have been shown to significantly diminish quality and activities of daily living(*13–16*). These non-motor symptoms also predict a steeper course of subsequent cognitive and motor decline, greater risk for freezing and falls and increased likelihood of ultimately developing PD-related dementia(*13, 17–21*). Moreover, because relatively few effective therapeutic options are available(*7, 8, 12, 13*), the early appearing mild cognitive and memory impairments of PD endure, progressively worsen and culminate in overall prevalence rates of more than 80%. The studies presented here provide further evidence for a recessive gene rat model–rats with knockout of PTEN-induced putative kinase 1 gene (Pink1-/-), as suitable for preclinical investigation of the biobehavioral factors that influence mild cognitive deficits in PD, including biological sex.

As in other neuropsychiatric conditions, significant sex differences have been shown to characterize key clinical features of PD (*22–25*). Most relevant to this study are consensus findings that link male sex to increased risk for mild cognitive deficits in PD and show that mild cognitive deficits appear earlier, with greater frequency and/or with greater severity in male compared to female patients with PD(*26–31*) In addition, sex differences have also been shown in the efficacies of an already limited pool of therapeutic options available to effectively treat these non-motor symptoms (*32–34*).

These and other findings have been critical drivers of a significant body of clinical research aimed at resolving the impacts of biological sex, sex hormones and hormone augmentation therapies on the pathophysiology and clinical management of cognitive and memory (and other) deficits in male and female patients with PD. While the work is ongoing, results to date are mixed and range mainly from promising to inconclusive (*35–39*). This is due in part to the limited availability of female patients with PD, population variance in clinical subtypes of disease and study-to-study differences in hormone augmentation regimens and other factors that impact patients’ health and hormone status. Importantly, these variables can be well-controlled in animal and especially rodent models of PD, several of which have been shown to be appropriate for exploring open questions about disease-related deficits in cognition and memory.

A number of mechanistically different rodent models of PD have been shown to recapitulate fundamental biobehavioral features of disease, including deficits in higher order information processing(*40–43*). The models that are most frequently used to explore cognition and memory are rats or mice in which pre-motor stages of PD-relevant pathology^1^ are induced by chemical agents, e.g., 6-hydroxydopamine (6-OHDA), 1-methyl-4-phenyl-1,2,3,6-tetrahydropyridine (MPTP), environmental toxins, e.g., rotenone, paraquat, or intracerebral injections of pre-formed α-synuclein fibrils (*43–50*). Behavioral testing in these and other models has revealed significant deficits in several higher-order processes, including spatial learning, behavioral flexibility, impulse control and memory. Several of these studies have further shown that deficits are more severe to exclusive in males and/or are sensitive to changes in sex hormone levels (*50–55*). While these elements of face validity are important, genetic rodent models of PD are also emerging as potentially useful in investigating cognitive and memory deficits in PD(*56, 57*). The broader benefits of these models include onsets of pathophysiological processes that are spontaneous and progressive and expressions of related deficits in motor and non-motor functions(*58–61*). Thus, they may be especially well suited for investigating temporarily dynamic aspects of mild cognitive impairments in PD, including the factors that influence their prodromal appearance, their rates of subsequent decline and their relationships to motor dysfunction. Resolving these questions has important implications for better understanding disease etiology, for identifying biomarkers and risk factors and for developing novel treatments that are potentially preventive, disease-modifying and beneficial to motor and non-motor signs.

In choosing genetic strains to address these issues, it may be important to consider that rats are widely viewed as the rodent of choice for studying cognition, memory and sex differences in these domains (*62–64*). Although limited in number, rat strains that are available for study include those that model autosomal dominant (e.g., LRKK2) or autosomal recessive (Pink1, Parkin, DJ-1) gene mutations that are clinically infrequent but of moderate to high risk/penetrance (*65, 66*). However, among the monogenic forms of PD related to these gene perturbations, patients where disease is caused by loss of PINK1 function have been shown to be most at risk for developing cognitive impairments(*67, 68*). This along with other findings, including recent evidence for vulnerability of neural correlates of cognition in Pink1-/- mice (*69*) have drawn the attention of our lab and others to Pink1-/- rats as potentially suitable models for cognition and memory loss in PD.

Loss-of-function PINK1 mutations are causally linked to early onset forms of PD (*70–72*) and are the second most common autosomal recessive gene disturbance leading this disease(*72*). In addition to construct validity, studies in Pink1-/- rats have shown that this strain also has face validity for a range of progressive, PD-relevant changes in mitochondrial function, α-synuclein accumulation and dysregulation/degeneration of monoamine, indolamine and cholinergic neurotransmitter systems (*65, 73–75*)(REFS).

These rats also develop somatic and orofacial motor deficits, deficits in the structure and intensity of ultrasonic vocalizations and behavioral expressions of anxiety and anhedonia that in some cases have been shown to be sex-specific (*76–78*). Recent studies have also shown that male Pink1-/- rats also express deficits in spatial cognition, i.e., complex route finding(*79*) and in performance of multiple object recognition-based memory tasks weeks to months before onsets of measurable deficits in, for example, hind limb strength or tapered balance beam traversal(*80*). The studies presented explore cognitive and memory loss in this strain further by longitudinally testing WT and Pink1-/- male and female rats on a series of object recognition-based memory tasks. The objectives of the work were to confirm previous evidence for the timing and severity of object recognition memory deficits in Pink1-/- males; to evaluate performance of female Pink1-/- for the first time in Novel Object Recognition (NOR), Novel Object Location (NOL) and Object in Place (OiP) tasks and to extend behavioral observations in Pink1-/- rats of both sex to constructs of episodic memory, which are among the first and the most frequently higher-order operations to be impaired in PD(*9, 10, 81, 82*). For the latter, male and female Pink1-/- rats and WT controls, were longitudinally tested on a modified version of the What, Where, When Episodic-like Memory task (WWW) in which olfactory cues were included as additional means that rats could use to distinguish sample objects. Mindful of the prevalence of hyposmia/anosmia in PD(*83, 84*), rats in all groups were also tested for olfactory discrimination and habituation, and estrous cycle tracking in female subjects was used to explore potential impacts of circulating hormones on all major behavioral outcome measures.

## 2. METHODS

### 2.1 Animal subjects

All procedures involving animals comply with the U.S. Public Health Service Guide for Care and Use of Laboratory Animals and with Stony Brook University’s institutional guidelines for laboratory animal use; all are also approved by Stony Brook University’s Institutional Animal Care and Use Committee (IACUC #227514; most recent date of approval is 6/23/2023). A total of 34 Long Evans rats were used: 10 were wild type (WT) females; 6 were WT males; 12 were *Pink1* knockout [*Pink1* -/-, (LE-Pink1^em1Sage-/-^) females; and 6 were *Pink1*-/- males. All rats were purchased from Inotiv, West Lafayette, IN, USA at 6-7 weeks of age and were double housed according to biological sex and genotype for the duration of the study. Home cages were standard translucent tub cages (Lab Products, Inc., Seaford, DE, USA) that contained ground corn cob bedding (Bed O’ Cobs, The Anderson Inc., Maumee, Ohio, USA), shredded nesting paper and enrichment objects (Nyla Bones, Nylabone, Neptune, NJ USA). Animals were housed under a 12-h non-reversed light-dark cycle with food (Purina PMI Lab Diet: ProLab RMH 3000) and water were available *ad libitum*.

### 2.2 Monitoring for Health and Estrous Cycle

All rats were handled at least once weekly and were weighed every 4-8 weeks. Estrous cycles in female rats were also tracked to establish cycle durations, regularity and stages on behavioral testing days. Beginning one week after their arrival, all females were vaginally lavaged with saline daily for two weeks. Thereafter, lavages were performed every 2-3 days. At ∼6 months of age, visual inspection of the vaginal opening replaced lavage as a less stressful method of identifying estrus, proestrus and diestrus phases of the estrous cycle (*85*). This was performed on behavioral testing days and intermittently in between.

### 2.3 Behavioral Testing

All testing took place in a core facility during rats’ subjective nights under ambient white light (∼ 260 lux). The core facility included a central holding room where animals were acclimated in home cages for ∼60 minutes. Subsequent behavioral testing took place in an adjacent 10-12 ft square, sound attenuated room equipped with an overhead digital camera (LogiTech webcams, San Jose CA) used to archive all trials. Beginning at 3 months of age, rats were tested every 2 months on a series of untrained, unrewarded behavioral paradigms measuring object recognition memories, affect, sensorimotor gating or motor function. Testing in females modeled a previous study conducted in male WT and Pink1-/- rats and extended analysis to episodic memory. Testing in males was designed to partially replicate earlier studies (NOR, NOL testing at 3 and 5 months), to pilot object memory tasks with longer retention contingencies (not reported) and to extend analyses in males to episodic memory. For rats of both sexes, the order of tasks was varied, and rats had 3-7 days off between tasks.

### 2.4. Testing Arenas, Apparatus and Procedures

#### 2.4.1 Object-based Memory Tasks

Object recognition-based memory tasks were performed in translucent, rectangular polypropylene testing arenas (32 in long, 19 in wide, 13 in high). One long wall of the arena was opaque and remaining walls had small, high contrast visual cues (stickers) fixed to their exteriors that were rearranged across testing months. During initial testing, the tops of the arenas were open and trials began by placing rats in opaque start cylinders located at the centers of the arenas. However, during testing at 5 months of age many female rats (WT and Pink1-/-) learned to climb the walls and escape. This negatively impacted their performance and that of other subjects being tested in parallel. Stopgap measures were tried to prevent this as testing at this age proceeded. Ultimately large plexiglass sheets were placed over the tops of the arenas and trials were initiated by placing rats directly in the arenas pointed away from objects. These adaptations were used for all rats for the remainder of the study, and data from female rats at the 5-month testing time point were excluded from analysis.

#### 2.4.2 Novel Object Recognition (NOR) Testing (Fig 1A)

Rats were given three 5-minute sample trials (S1, S2, S3) each separated by 1 hour and one 5-minute test trial (TT) 1.5 hours after the third sample trial. During the sample trials, rats were free to explore two identical objects that were placed in adjacent corners of the arena, at least 4 in from the arena walls. During the test trial, objects were in the same corner locations as the sample trials. However, one object was from the sample trials (familiar object) and the second object was one the rats had never encountered (novel object). Intertrial intervals were spent in home cages in the adjacent holding room.

**Figure 1.**
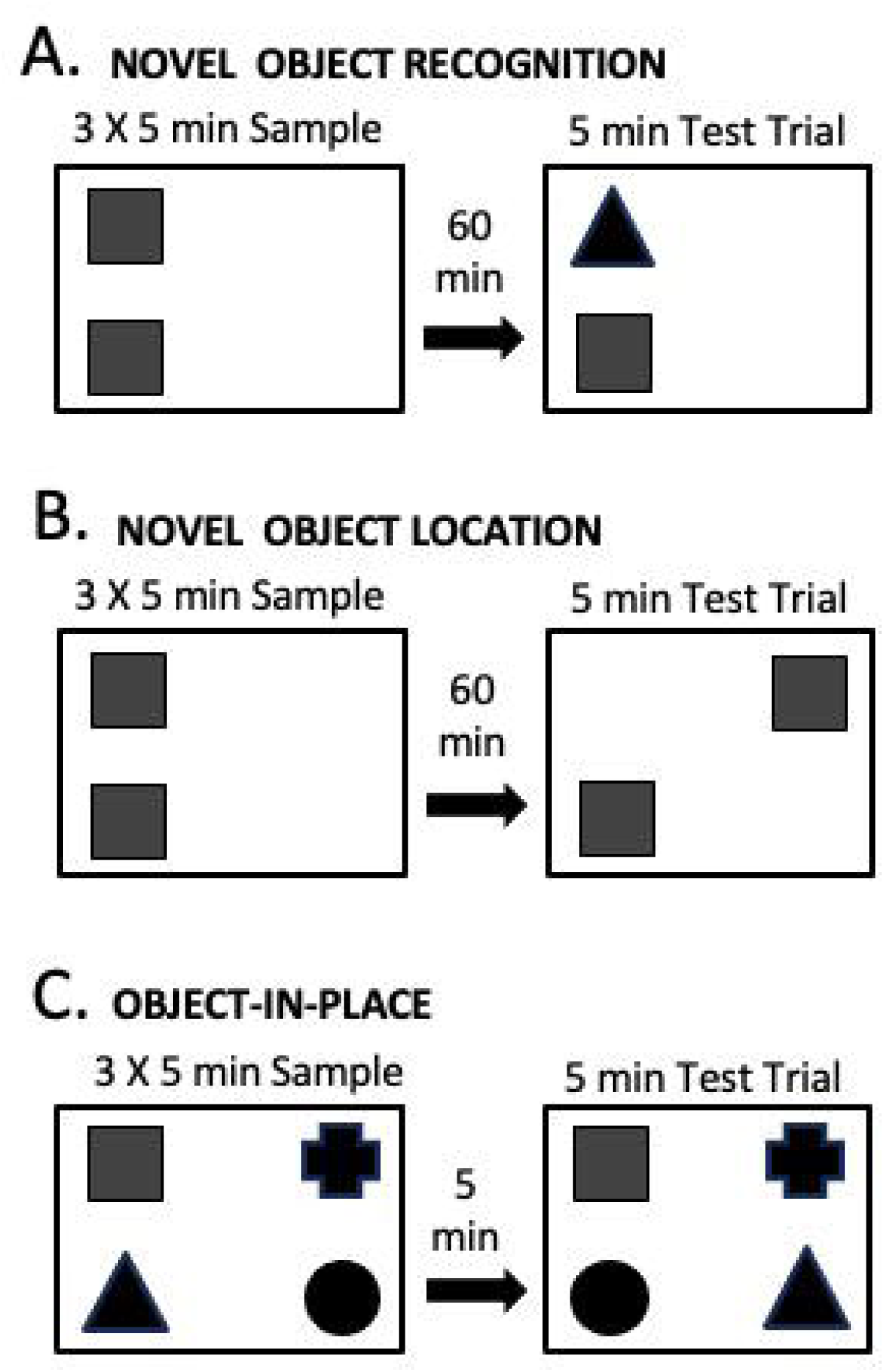
Schematic diagrams showing testing arena setups and trial structures for behavioral testing. Panel A shows that for the Novel Object Recognition paradigm, rats are given three sample trials of 5 minutes’ durations to view two identical sample objects; during the subsequent test trial, one of the objects (familiar) is replaced with a new one (novel). Panel B shows that for the Novel Object Location paradigm, rats are given three sample trials of 5 minute’s duration to view two identical sample objects; during the subsequent test trial, one of the objects is presented in the same place (familiar) and one is presented in a new (novel) location. Panel C shows that for the Object-in-Place paradigm, rats are given three sample trials of 5 minute’s duration to view 4 distinct sample objects; during the subsequent test trial, the positions of two of the objects are the same (familiar) and the positions of the other two are swapped with each other (novel).

#### 2.4.3 Novel Object Location (NOL)Testing (Fig 1B)

Rats were given three 5-minute sample trials (S1, S2, S3) each separated by 1 hour and one 5-minute test trial (TT) one hour after the third sample trial. During the sample trials, rats were free to explore two identical objects. The locations of the objects varied at each age tested but were always at least 4 in from the arena walls. The test trials used the same objects that were presented during the sample trials. One was in the same place that it had been during the sample trials (familiar location), and the other was in a place that neither object had been previously located (novel location). Intertrial intervals were spent in home cages in the adjacent holding room.

#### 2.4.4

(S1, S2, S3) that were separated by 5 minutes and a single 3-minute test trial (TT) that began 5 minutes after the third sample trial. During the sample trials, rats were free to explore 4 unique objects that were placed in each of the four corners of the arena, at least 4 in from the arena walls. During the test trial, rats were free to explore the same four objects. However, two of the objects remained in their original sample trial locations, and the other two objects switched locations. Intertrial intervals were spent in home cages in the adjacent holding room.

#### 2.4.5 What Where When Episodic-Like Memory (WWW)Testing (Fig 2A)

Rats were given two unique 5-minute sample trials (S1, S2) and one 5-minute test trial (TT) with all trials separated by 50-minute intertrial intervals. During the sample trials (S1, S2), rats were presented with one of two quadruplicate sets of objects. For S1 trials, objects were triangularly arrayed and during S2 trials the objects were placed near the arena corners to form squares. During test trials (TT) the objects were arranged in the S2 square configuration. However, two objects were those from S2 (recent familiar, RF) and two were from S1 (old familiar, OF). Both of the S2 objects occupied previously held positions. For the S1 objects, one occupied a previously held location (old familiar stationary, OFS) and the other was presented in a new location (old familiar displaced, OFD).

**Figure 2.**
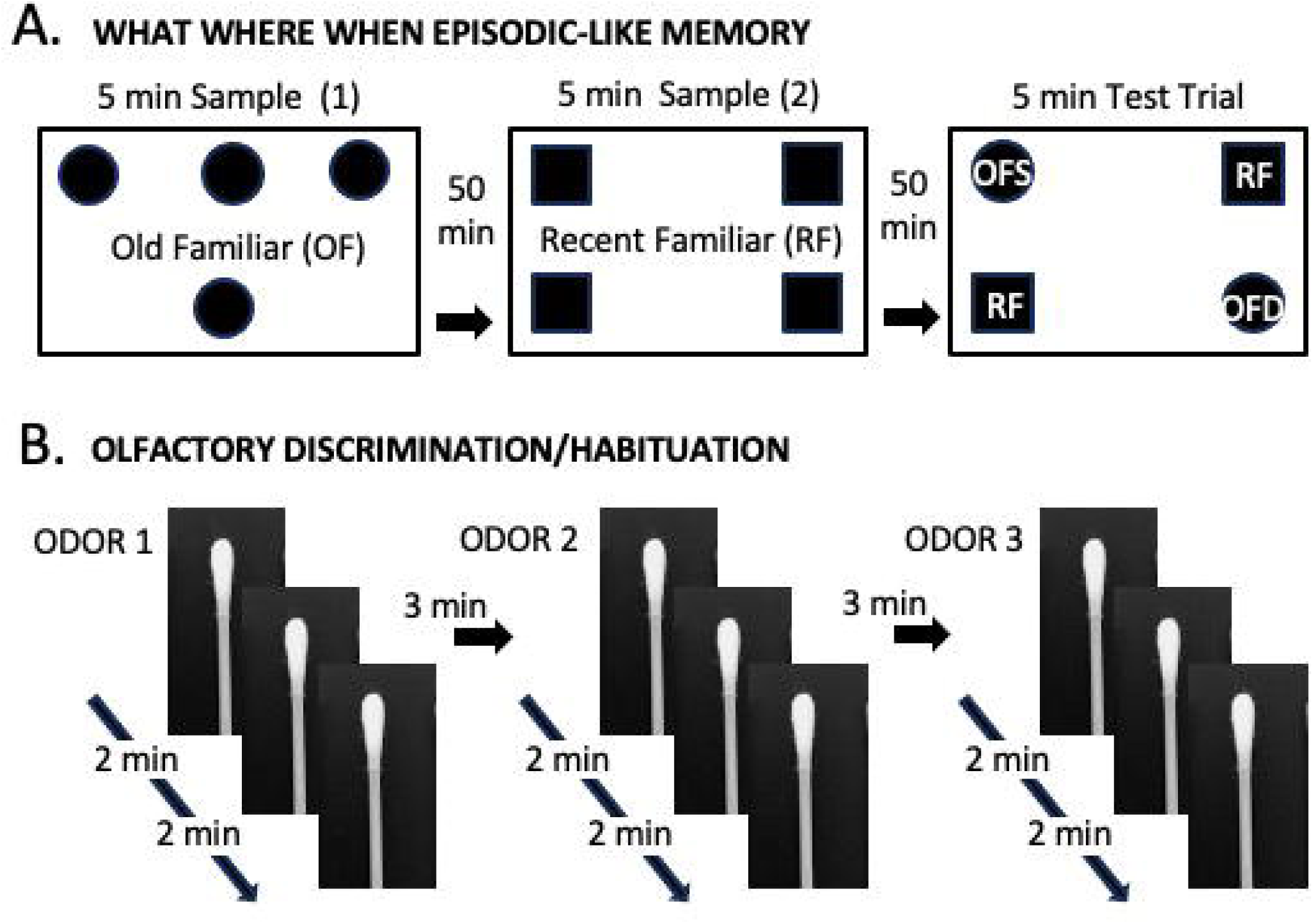
Schematic diagrams showing testing arena setups and/or trial structures for behavioral testing. Panel A shows that for the What, Where, When Episodic-Like Memory paradigm, rats are given two consecutive sample trials of 5 minutes’ durations. In the first, a quadruplicate set of sample objects are arrayed in a triangle; in the second sample trial a different quadruplicate set of objects are arranged in a square. During the subsequent test trial, two objects from the second sample are presented in previously occupied positions (Recent Familiar; RF). Two objects from the first trial are also presented; one is in a previously occupied position (Old Familiar Stationary; OFS), and one is in a new position (Old Familiar Displaced; OFD). Panel B shows the sequential presentations of three distinct non-social odor solutions using soaked cotton-tipped applicators. Odor discrimination and habituation are measured by presenting each odor 3 times before switching to the next odor solution.

#### 2.4.6 Sample and Test Trial Objects and Object Locations

For NOR, NOL, and OIP testing at 3 and 7 months and for NOR and NOL testing in males at 5 months, the objects used were largely the same as those used in a previous study of WT and Pink1-/- male rats at these same ages (*80*). NOR, NOL and OIP testing at 12 months also used previously used objects, but in different pairings. In all cases, objects serving as familiar (sample) or novel stimuli were similar in overall size and/or shape and differed along limited dimensions, e.g., of color/contrast, composite material, e.g., glass, plastic, metal and/or surface features, e.g., smooth, grooved. Objects used for NOL testing also had depressions, handles or other features to encourage exploration. Objects serving as novel or familiar stimuli (NOR), arena zones that served as sample and novel locations (NOL) and arena corners where objects were switched or remained stationary (OIP) were all counterbalanced across subsets of rats in all groups.

For WWW testing, the quadruplicate sets of objects used were also similar to one another in overall dimensions and were made of the same materials (plastic, metal and/or glass) but differed in shape, color/opacity and/or surface texture. The objects were also hollow and had small holes present or drilled in their tops to allow access to distinguishable odorant stimuli that were placed inside. Odorants were matched by categories of spice (fennel vs. anise), savory (canned dog food vs. mild cheddar cheese) or sweet (milk chocolate chips vs butterscotch chips) and were weighed to ensure that equal amounts were present in each of the four quadruplicate objects. The object/odor sets used in S1 (old familiar objects) or S2 (recent familiar objects) trials were counterbalanced across subsets of animals in all groups. The exception to this was testing male rats at 12 months of age where the same object sets were inadvertently used in S1 and S2 trials for all subjects.

#### 2.4.7 Olfactory Task

A non-social odor discrimination and habitation task modeled on Yang and Crawley 2009 (*86*), was performed in clean home cages that contained no bedding. At the center of one end of the enclosure, a clamp was fixed to the floor to hold (and to rapidly replace) single 4-inch wooden cotton-tipped applicators upright (cotton tip up). A clear plexiglass sheet was placed over the top of the arena to prevent escape. Rats were tested on this paradigm one time at 5-6 months of age.

#### 2.4.8 Odor Discrimination/Habituation Testing (*Fig 2B*)

Rats were placed in the empty arena and given 3 minutes to acclimate. Next, a dry cotton-tipped applicator was placed in the holding clamp and rats had another 3 minutes to acclimate to its presence. Finally, the dry applicator was removed and sequentially replaced with a series of fresh applicators with cotton tips that were dipped in one of three distinct odorant solutions. Each odorant was presented in three successive 2-minute trials that were separated by 3-minute intertrial intervals. The arenas and applicator clamps were wiped with dry paper towels between odor presentations and were cleaned with 70% EtOH before and after testing different animal subjects.

#### 2.4.9 Odorant Solutions

The three odorant solutions used were: 1. Distilled water; 2. Banana extract (McCormick, Hunt Valley, MD) diluted 1:100 in distilled water; and 3. Almond extract (McCormick, Hunt Valley, MD) diluted 1:100 in distilled water.

### 2.5 Data Analysis

#### 2.5.1 Estrous Cycle Determination

Vaginal cytology samples were evaluated using light microscopy and a combination of brightfield and differential interference contrast illumination. Estrous cycle stages were cytologically identified as follows:

⍰ Proestrus: round, nucleated epithelial cells are most abundant.
⍰ Estrus: cornified and anucleated epithelial cells are most abundant.
⍰ Diestrus: leukocytes are most abundant.

Estrous cycle stage was also evaluated using visual inspections of the vaginal opening as follows:

⍰ Estrus or Proestrus: vaginal opening is wide to gaping; surrounding tissues appear pink in color.
⍰ Diestrus: vaginal opening is narrow to closed; surrounding tissues appear white to bluish in color.

#### 2.5.2 Measurements of Object Exploration/Investigation

Object exploration during sample trials was assessed using automated tracking software (EthoVision XT 17.0) and was defined as the amount of time that nose point tracking out the locations of rats’ snouts within 2 cm of the objects. During test trials, object exploration was analyzed from digitally recorded trials by trained observers using event capture software [Behavioral Observation Research Interactive Software (BORIS) version 7.8.2, open access] and was defined as amounts of time rats spent actively sniffing or whisking the objects.

#### 2.5.3 Calculation of Discrimination Indices (DI)

Object memories were evaluated during test trials by calculating discrimination indices. For NOR and NOL, these indices represented differences in the amounts of time (sec) rats spent exploring objects that were novel (NOV) vs. familiar (FAM) or that were located in novel (NOV) vs. familiar (FAM) locations. For both tasks, DI was calculated using the following formula:

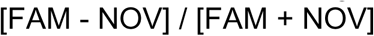

For OIP, discrimination indices reflected differences in the amounts of time rats spent with objects that were located in swapped or displaced (DSPL) vs. stationary/previous locations (STA) and were calculated using the following formula:

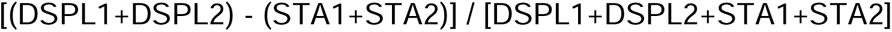

For the WWW task, discrimination indices reflected episodic memory modalities of “what”, “where” and ‘when” based on differences in the amounts of time rats spent exploring objects that were defined as recent familiar (RF, presented in Sample Trial 2) as old familiar (OF, presented in Sample Trial 1), or as old familiar displaced (OFD, from Sample Trial 1 but in a new location) or as old familiar stationary (OFS, from Sample Trial 1 and located in a previously occupied position). These indices were calculated by the following formulas:

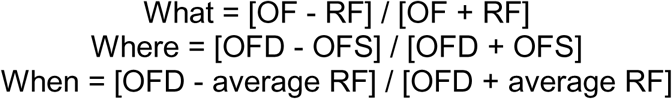

These equations constrain DI values from 1 to −1, and yield DI values of zero when object pairs in question are investigated equally, i.e., no discrimination.

#### 2.5.4 Olfactory Discrimination/Habituation

Measures of olfactory discrimination and habitation were analyzed from digitally recorded trials by trained observers using event capture software (BORIS). These analyses scored the amounts of times rats spent actively sniffing cotton tipped applicators during three sets of three trials each where applicators were sequentially soaked in the different odorant solutions.

### 2.6 Statistics

Data from small numbers of individual rats from all groups were excluded from the analysis if they froze, failed to minimally investigate objects (5 sec) or if any contamination of a sample object, e.g., urine droplet, was detected. Finally, the Grubbs’ test was used to identify outliers in all data sets; this test revealed very small numbers of data points across all tasks and groups that were removed from the analysis.

Statistical analyses were performed using IBM SPSS, Version 25 (SPSS, Inc., Chicago, IL, USA). All data were first assessed for descriptive statistics, including Levine’s F-test for equality of variance. Body weights and discrimination indices (DI) for NOR, NOL and OIP tasks were compared across groups using one-way analyses of variance (ANOVA). Outcome measures for olfactory discrimination/habituation and episodic memory tasks, were evaluated using ANOVAs with repeated-measures designs. For these, Mauchly’s test for sphericity of the covariance matrix was applied, degrees of freedom were adjusted as needed using the Huynh-Feldt epsilon. Where allowed, follow-up pairwise comparisons (one way ANOVAs) were used to identify significant group differences. To assess estrous cycle impacts, ANOVAs and repeated measures ANOVAs were applied as above albeit with ‘groups’ re-defined and WT or Pink1-/- females that were in proestrus or estrus vs. diestrus on the day of testing. Discrimination index data were also evaluated using within-groups one sample t-tests to identify DI values as significantly different–or not, from zero and relationships between DI and total sample trial object exploration times were assessed by applying Pearson’s regression analyses within groups. Effect sizes were also assessed by calculating Cohen’s d (*d*) for t-tests or eta squared (η^2^) for ANOVAs.

## 3. RESULTS

### 3.1 Body Weights

The body weights of wildtype (WT) male and female rats were commensurate with age, increased steadily over time and were markedly higher in males compared to females at every point evaluated (Fig 3). At 3 months of age, weights in the Pink 1-/- groups were also noticeably greater than those of sex-matched WT controls. However, at 7 months these group differences were diminished and by the 12-month time point they were smaller still to negligible. All of these observations were supported statistically. First, a repeated measures ANOVA that included all four groups and all three timepoints identified significant main effects of Age [F_(1.28,3.84)_ = 235.49, p < 0.001; η^2^ = 0.89 (large)], significant main effects of Group [F_(3,29)_ = 332.16, p < 0.001; η^2^ = 0.97 (large)] and significant interactions between these variables [F_(3.84,37.07)_ = 20.25, p < 0.001; η^2^ = 0.68 (large)]. Follow pairwise comparisons further showed that weights in WT males were significantly greater than WT females at all ages (p < 0.001); that weights in Pink1-/- males were significantly greater than WT males at 3 (p < 0.001, FIg 3A) but not at 7 or 12 months of age; and that weights in Pink1-/- females were significantly greater than WT females at 3 and 7 but not 12 months of age (p < 0.001, Fig 3B).

**Figure 3.**
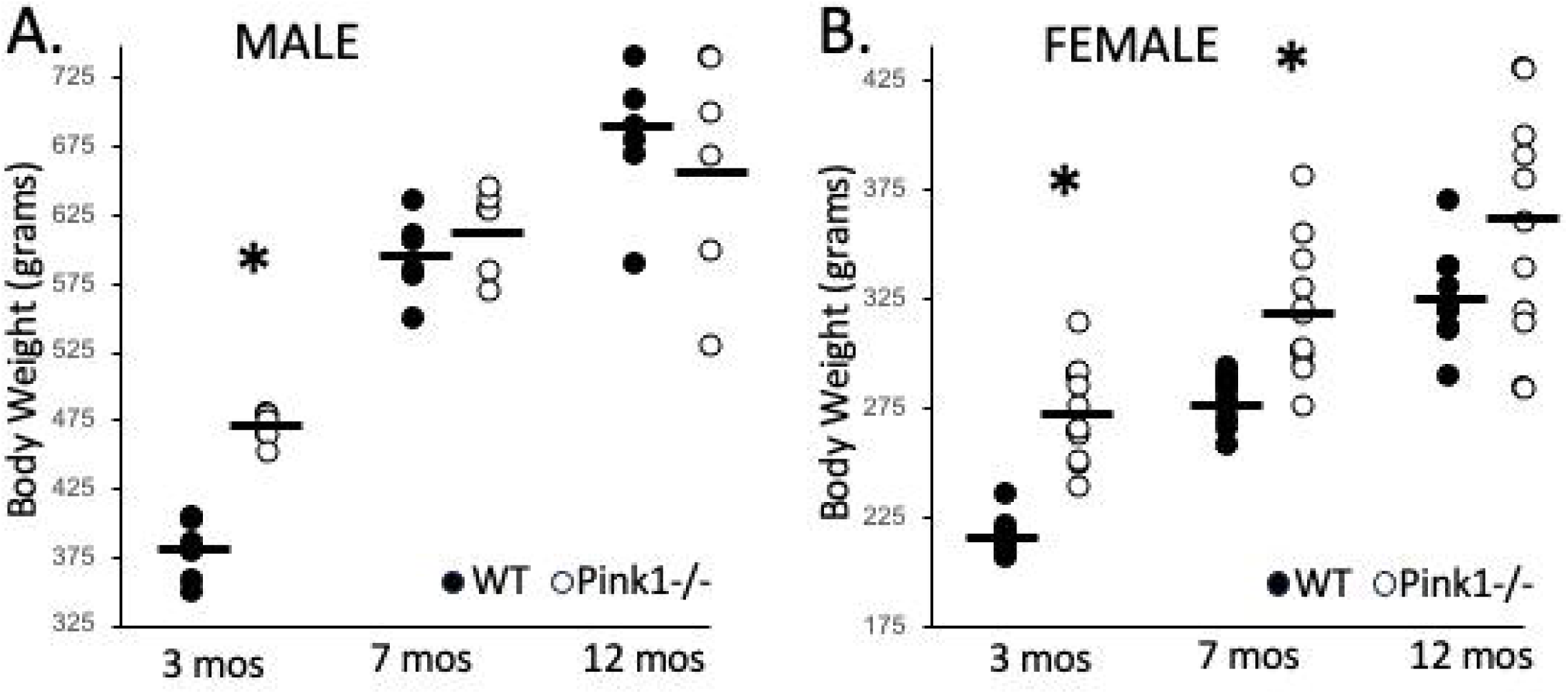
Scatter plots showing the body weights in grams of male (A) and female (B) wild type (WT, black circles) and Pink1 knockout (Pink1-/-, open circles) rats. Both groups of male rats weighed roughly twice as much as the corresponding groups of female rats. Among males, the body weights of the Pink1-/- rats were significantly greater than those WT at 3 months of age but were not significantly different than the controls at 7 or 12 months of age. For the females, the body weights of the Pink1-/- rats were significantly greater than those WT at 3 and 7 months of age but were not significantly different than the controls at 12 months old.

### 3.2 Estrous Cycles

Daily analyses of vaginal cytology samples conducted over a two week period established regular 4-day estrous cycling in all subjects in both the WT and Pink1-/- female groups prior to the onset of behavioral testing. Cytological sampling or visual inspections thereafter identified estrous cycle stage on testing days and intermittently in between. There were no indications that estrous cycles were impacted by genotype, were interrupted by behavioral testing or significantly lengthened as animals aged. The numbers of rats that were in estrus or proestrus vs. diestrus on testing days are presented separately for each task in Table 1.

**Table 1.** Estrous cycle stages were tracked on days of behavioral testing. The numbers of Wild Type and Pink1 knockout (Pink1-/-) female rats that were in estrus or proestrus vs. diestrus during testing for the Novel Object Recognition (NOR), Novel Object Location (NOL), Object-in-Place (OIP), What, Where, When Episodic-like memory (WWW) or Olfactory discrimination and habituation (OLF) tasks are shown for each age that testing took place.

| **NOR Task** | **3 Months of age** | **7 Months of age** | **12 Months of age** |
| --- | --- | --- | --- |
| *Wild Type* | Estrus/Proestrus, n = 3<br>Diestrus, n = 7 | Estrus/Proestrus, n = 9<br>Diestrus, n = 1 | Estrus/Proestrus, n = 8<br>Diestrus, n = 2 |
| *Pink1<sup>-/-</sup>* | Estrus/Proestrus, n = 7<br>Diestrus, n = 5 | Estrus/Proestrus, n = 7<br>Diestrus, n = 5 | Estrus/Proestrus, n = 4<br>Diestrus, n = 7 |
| **NOL Task** | **3 Months of age** | **8 Months of age** | **12 Months of age** |
| *Wild Type* | Estrus/Proestrus, n = 8<br>Diestrus, n = 2 | Estrus/Proestrus, n = 7<br>Diestrus, n = 3 | Estrus/Proestrus, n = 10<br>Diestrus, n = 0 |
| *Pink1<sup>-/-</sup>* | Estrus/Proestrus, n = 8<br>Diestrus, n = 4 | Estrus/Proestrus, n = 3<br>Diestrus, n = 9 | Estrus/Proestrus, n = 10<br>Diestrus, n = 2 |
| **OIP Task** | **3 Months of age** | **8 Months of age** | **12 Months of age** |
| *Wild Type* | Estrus/Proestrus, n = 5<br>Diestrus, n = 5 | Estrus/Proestrus, n = 10<br>Diestrus, n = 0 | Estrus/Proestrus, n = 4<br>Diestrus, n = 6 |
| *Pink1<sup>-/-</sup>* | Estrus/Proestrus, n = 4<br>Diestrus, n = 8 | Estrus/Proestrus, n = 4<br>Diestrus, n = 8 | Estrus/Proestrus, n = 5<br>Diestrus, n = 6 |
| **WWW Task** | **3 Months of age** | **8 Months of age** | **12 Months of age** |
| *Wild Type* | Estrus/Proestrus, n = 4<br>Diestrus, n = 6 | Estrus/Proestrus, n = 8<br>Diestrus, n = 2 | Estrus/Proestrus, n = 5<br>Diestrus, n = 5 |
| *Pink1<sup>-/-</sup>* | Estrus/Proestrus, n = 5<br>Diestrus, n = 7 | Estrus/Proestrus, n = 7<br>Diestrus, n = 5 | Estrus/Proestrus, n = 9<br>Diestrus, n = 3 |
| **OLF Task** | **5-6 Months of age** |  |  |
| *Wild Type* | Estrus/Proestrus, n = 8<br>Diestrus, n = 2 |  |  |
| *Pink1<sup>-/-</sup>* | Estrus/Proestrus, n = 6<br>Diestrus, n = 6 |  |  |

### 3.3 Object Recognition Memories: Replication Studies in Males

Wild type and Pink1-/- male rats were tested on NOR and NOL tasks at 3 and 5 months of age. The results obtained replicated findings from a previous study of object recognition memories in male Pink1-/- rats (REF). Initially, at 3 months of age, WT and Pink 1-/- male rats showed similarly strong discrimination of novel compared to familiar objects [mean discrimination index (DI) values of 0.24 and 0.48, respectively] and similarly poor discrimination of objects based on spatial location (DI’s for NOL: WT = 0.09; Pink1-/- = - 0.08, Fig 4A). Analyses of variance confirmed that there were no significant group differences in mean DI values for either task (NOR: η^2^ = 0.29; NOL: η^2^ = 0.14). One-sided t-tests further showed that for both groups DI values were significantly greater than zero for NOR testing [WT: *t*(5) =3.41, p = 0.02, *d* = 1.75; Pink1-/-: *t*(5) = 4.93, p = 0.004, *d* = 0.36] and were not significantly different than zero for NOL testing (WT: *d* = 0.36; Pink1-/-: *d* = 0.33). At 5 months of age, WT males showed robust NOR and NOL discrimination (DI’s 0.44 and 0.24, respectively, Fig 4D) while Pink1-/- males showed poor performance in both tasks (NOR: DI = 0.16; NOL: DI = −0.08, Fig 4D). Analyses of variance confirmed that group differences in DI were significant for both the NOR [F_(1,10)_ = 14.78, p = 0.004; η^2^ = 0.622] and the NOL task [F_(1,10)_ = 6.87, p = 0.03; η^2^ = 0.41]. One sided t-tests further showed that DIs for both tasks were significantly greater than zero in WT rats [NOR: *t*(5)= 3.92, p = 0.01, *d* = 1.13; NOL *t*(5)= 4.04, p = 0.01, *d* = 0.29]. However, in the Pink1-/- cohort, mean DIs were only significantly greater than zero for NOR [NOR: t(5) = 3.90, *p* = 1.13, *d* = 0.102; NOL: *d* = 0.29).

**Figure 4.**
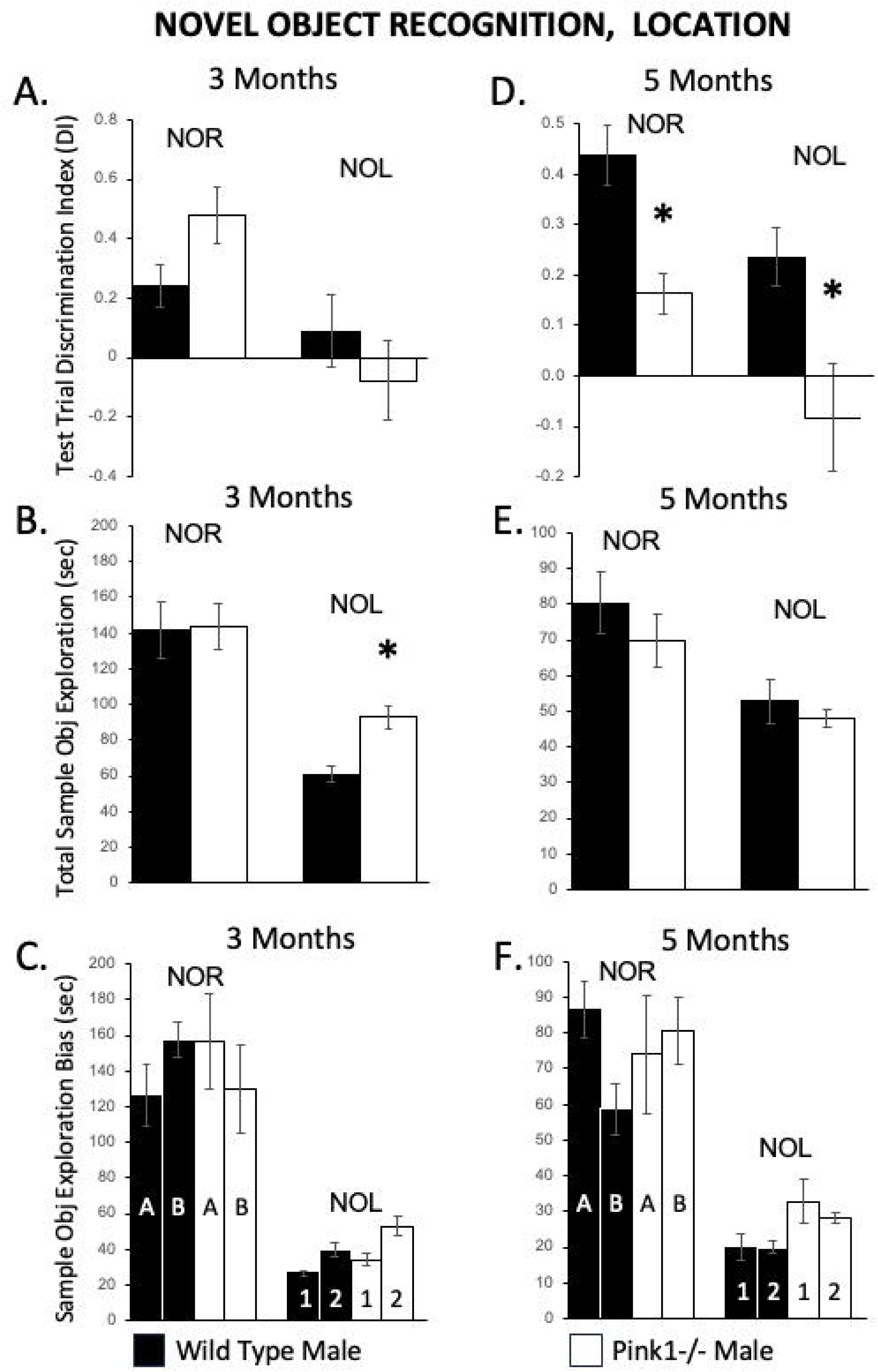
Bar graphs summarizing test trial (A, D) and sample trial (B, C, E, F) data for Novel Object Recognition (NOR) and Novel Object Location (NOL) testing in wild-type (WT; black bars) and Pink1 knockout (Pink1-/-; white bars) male rats. Rats were tested at 3 (left-hand column) and 5 months of age (right-hand column). At 3 months of age, discrimination indices (DI) in Pink1-/- rats and WT controls were similarly robust for NOR and similarly weak for NOL (A). However, at 5 months of age DI values for both NOR and NOL were significantly (*) lower in Pink1-/- compared to WT rats (D). Total sample trial object exploration times [B, E in seconds (sec)] were mostly similar in Pink1-/- and WT rats. The only exception was that at 3 months of age, the Pink1-/- rats spent significantly longer (*) than WT rats investigating NOL sample objects (B). There were no significant differences noted within or across groups in total sample object exploration times for data that were stratified by counterbalanced objects (bars marked A or B) that were used as familiar vs. novel stimuli in NOR testing and no evidence of spatial bias in the amounts of times rats spent with objects placed in each of the two locations (bars marked 1 or 2) in NOL testing.

Within and across groups ANOVAs or repeated measures ANOVAs were also performed on sample trial exploration data to probe for potentially confounding effects of sample object exposure or for object or object location bias on test trial measures of discrimination. Comparisons of total sample object exploration (Fig 4B, D) identified only one significant group difference in this measure: NOL testing at 3 months of age [F_(1,10)_ = 16.67, p = 0.002; η^2^ = 0.63, Fig 4B]. This difference was driven by Pink1-/- rats spending roughly 50% more time (∼90 vs. 60 sec) than WT males exploring the pair of sample objects. No other comparisons revealed significant differences in rats’ explorations and calculated effect sizes that were consistently small (η^2^ =0.001-0.081, Fig 4 B, E). Further, regression analyses identified only one case of a significant correlation between total sample object exploration time and test trial DI; this was for NOR testing in 5-month-old WT rats (R^2^= 0.87, *p* = 0.007). All other correlations were non-significant (R^2^ = 0.00-0.52, *p* = 0.10-0.99). Finally, no significant differences were seen in either group in the total amounts of time rats spent with counterbalanced objects used as novel vs. familiar stimuli (NOR 3 mos: WT: η^2^ = 0.05; Pink1-/-: η^2^ = 0.22; NOR 5 mos: WT: η^2^ = 0.10; Pink1-/-: η^2^ = 0.46, Fig 4C,F left) or with sample objects in a given location (NOL 3 mos: WT: η^2^ = 0.05; Pink1-/-: η^2^ = 0.41; NOL 5 mos: WT: η^2^ = 0.10; Pink1-/-: η^2^ = 0.57, Fig 4C,F right).

### 3.4 Object Recognition Memories: New Studies in Females

#### 3.4.1 Novel Object Recognition

In NOR testing, WT and Pink 1-/- female rats both spent at least twice as long exploring novel compared to familiar objects. At 3, 7 and 12 months of age, mean DI values for the WT group were 0.50, 0.30 and 0.42, respectively, and corresponding DIs in the Pink1-/- cohort were 0.31, 0.29 and 0.42 (Fig 5A). Although analyses of variance identified a significant group difference in DI values at 3 mos of age [F_(1,19)_ = 4.65, p = 0.04; η^2^ = 0.20, FIg 5A], one-sided t-tests confirmed that DI values in both groups were highly robust, and significantly greater than zero [3 mos WT: *t*(9) = 6.74, p < 0.001, *d* = 2.13; 3 mos Pink1-/-: *t*(10) = 6.43, p < 0.001, *d* = 1.94]. Testing at 7 and 12 months of age, revealed no significant group differences in NOR DIs, calculated effect sizes were small (η^2^= 0.000 for both time points) and one-η^2^ sided t-tests confirmed that DI values in both groups were significantly greater than zero [7 mos WT: t(9) = 2.95, p = 0.02, *d* = 0.36; 7 mos Pink1-/-: *t*(11) = 3.55, p = 0.005, *d* = 0.82; 12 mos WT: *t*(9) = 4.90, p < 0.001, *d* = 1.55; 12 mos Pink1-/-: *t*(10) = 6.22, p < 0.001, *d* = 01.88]. Finally, although variably powered, within groups comparisons (ANOVA) performed on mean DI data that were stratified by rats’ estrous cycle stages on the day of testing revealed no significant effects of estrous cycle on NOR DIs for either group at any age (η^2^ = 0.04-0.11). The variable and in some cases poorly balanced numbers of WT and Pink1-/- subjects that were in estrous or proestrus vs. diestrus that were compared are shown in Table 1. These numbers argue that caution should be applied in drawing firm conclusions about estrous cycle impacts from these data.

**Figure 5.**
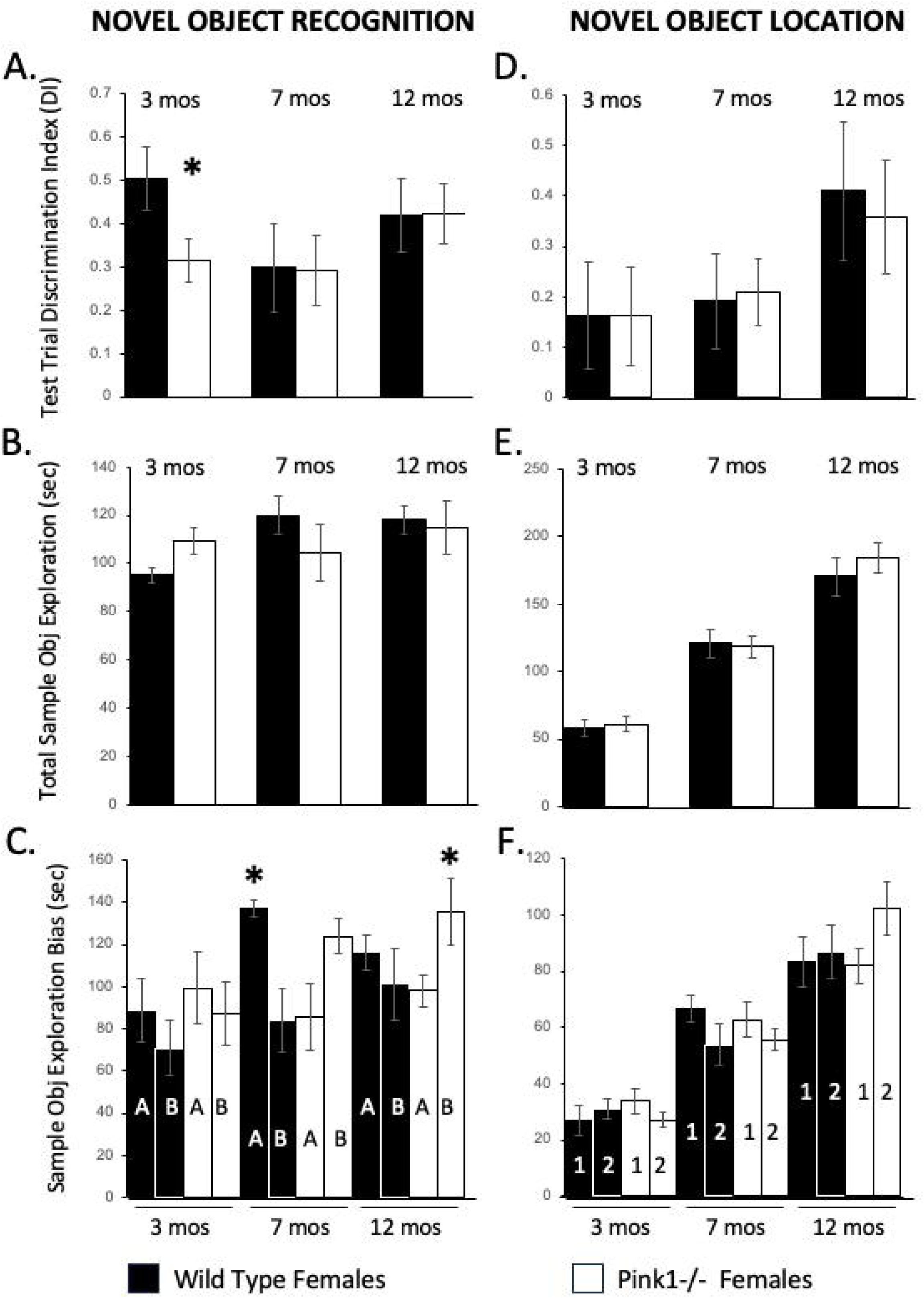
Bar graphs summarizing test trial (A, D) and sample trial (B, C, E, F) data for Novel Object Recognition (NOR, left hand column) and Novel Object Location (NOL, right hand column) testing in wild-type (WT; black bars) and Pink1 knockout (Pink1-/-; white bars) female rats. Data from testing at 3, 5 and 7 months of age are shown. The only instance where DI values were significantly lower in Pink1-/- compared to WT rats (*) was in NOR testing at 3 months (A); in this instance, DI values in both groups were robust, but were unusually high for the WT females. In all other testing, no significant group differences were observed. Analyses of total times spent investigating objects [in seconds (sec)] during sample trials also revealed no significant group differences for either the NOR (B) or NOL (E) task. There were two cases noted where significant group differences (*) were noted in total sample object exploration times for data that were stratified by counterbalanced objects (bars marked A or B) that were used as familiar vs. novel stimuli in NOR testing. At 7 months of age, WT rats spent more sample trial time exploring when Object A was the familiar stimulus and at 12 months of age Pink1-/- rats spent more sample trial time exploring when Object B was the familiar stimulus. There was no evidence of spatial bias in NOL testing at any age in the amounts of time rats of either group spent with objects placed in each of the two locations (bars marked 1 or 2).

Statistical evaluations of sample trial object exploration behaviors were also carried out. These comparisons found no significant group differences in total sample exploration times and calculated effect sizes that were small (η^2^ = 0.05-0.0030, Fig 5B). Regression analyses also found no significant correlations between total sample object exploration times and test trial performance for either group at any age (R^2^= 0.003-0.23, *p* = 0.24-0.87. Finally, within groups ANOVAs found only isolated cases of significant differences in the total amounts of time that WT or Pink1-/- females spent with the one or the other counterbalanced sample objects (Fig 5C). These were limited to 7 month old WT females [F_(1,8)_ = 11.90, p = 0.009; η^2^ = 0.60] and to 12 month old Pink1-/- rats [F_(1,9)_ = 6.77, p = 0.03; η^2^ = 0.43, Fig 5C]. In both bases, differences were driven by rats spending about 45 sec more investigating one of the counterbalanced sample objects compared to the other. Importantly, however, follow-up analyses of variance that tested whether these biases affected test trial performance found no significant differences in mean DI values of either WT (η^2^ = 0.14) or Pink1-/- rats (η^2^ = 0.14) that used one or the other objects as the familiar sample.

#### 3.4.2 Novel Object Location

At 3 months of age female WT and Pink 1-/- rats both showed poor discrimination of objects based on spatial location (WT: DI = 0.16; Pink1-/-: DI = 0.16, Fig 5D). Analysis of variance confirmed that there were no significant group differences in these NOL DI values and calculated effect size were small (η^2^ = 0.000). One-sided t-tests also showed that neither DI value was significantly different from zero (WT: *d* = 0.49; Pink1-/-: *d* = 0.48). However, at 7 and at 12 months of age, spatial discrimination was robust in both groups (7 mos: WT DI = 0.19, Pink1-/- DI = 0.21; 12 mons: WT DI = 0.41, Pink1-/- DI = 0.39, Fig 5D). For both groups and both timepoints, analyses of variance identified no significant group differences in DI measures, calculated effects sizes were small (7 mos η^2^ = 0.002; 12 mos η^2^ = 0.005) and one-sided t-tests identified all DI values as significantly to near significantly greater than zero [7 mos WT: *t*(7) = 2.04, p = 0.08, *d* = 0.72; 7 mos Pink1-/-: *t*(11) = 3.18, p = 0.009, *d* = 0.92; 12 mos WT: *t*(7) = 2.99, p = 0.02, *d* = 1.06; 12 mos Pink1-/-: *t*(10) = 3.20, p = 0.009, *d* = 0.97]. Analyses of variance performed on group mean DI data that were stratified by estrous cycle stage found no significant effects of high vs. low circulating ovarian hormone levels on DI at any testing age (η^2^ = 0.06-0.22). The numbers of WT and Pink1-/- subjects in estrous or proestrus vs. diestrus that were compared in these analyses (Table1) indicate that several of these comparisons are significantly underpowered.

Analyses of sample trial object exploration behaviors found no significant group differences in total sample object exploration times for either group at any age (η^2^ = 0.07-0.11, Fig 5E). Further, although regression analyses identified a significant positive correlation between sample object exploration and DI in WT rats at 12 months of age (R^2^= 0.63, *p* = 0.02), no other correlations were significant (R^2^= 0.02-0.32, *p* = 0.06-0.76). Finally, repeated measures ANOVAS found no evidence for significant spatial bias, i.e., differential exploration of one or the other sample object, for either group at any age (WT: η^2^ = 0.01-0.29; Pink1-/-: η^2^ = 0.10-0.25, Fig 5F).

#### 3.4.3 Object-in-Place

Wild type and Pink 1-/- female rats showed robust discrimination of objects based on relative positioning at 3, 7 and 12 months of age (Fig 6A). For WT females, DI values ranged from 0.30-0.37 and for the Pink1-/- cohort the corresponding range was 0.29-0.38(Fig 6A). Analyses of variance found no significant group differences in OIP DI measures at any age, and calculated effect sizes were low (η^2^ = 0.001-0.05). One-sided t-tests further showed that all DI values for both groups were significantly greater than zero [3 mos WT: *t*(8) = 4.73, p < 0.001, *d* = 1.58; 7 mos WT: *t*(8) = 2.30, p = 0.05, *d* = 0.77; 12 mos WT: *t*(8) = 2.44, p = 0.04, *d* = 0.81; 3 mos Pink1-/-: *t*(9) = 6.15, p < 0.001, *d* = 1.94; 7 mos Pink1-/-: *t*(8) = 2.51, p = 0.04, *d* = 0.84; 12 mos Pink1-/-: *t*(10) = 3.55, p = 0.005, *d* = 1.07]. Analyses of variance performed on mean DI data from WT and Pink1-/- groups that were stratified by estrous cycle stage found no significant effects of high vs. low circulating ovarian hormone levels on DI at any testing age (η^2^ = 0.02-0.37). It is important to note that the numbers of WT and Pink1-/- subjects in estrous or proestrus vs. diestrus that were included in these analyses (Table 1) were evenly divided for some comparisons, but highly unbalanced for others.

**Figure 6.**
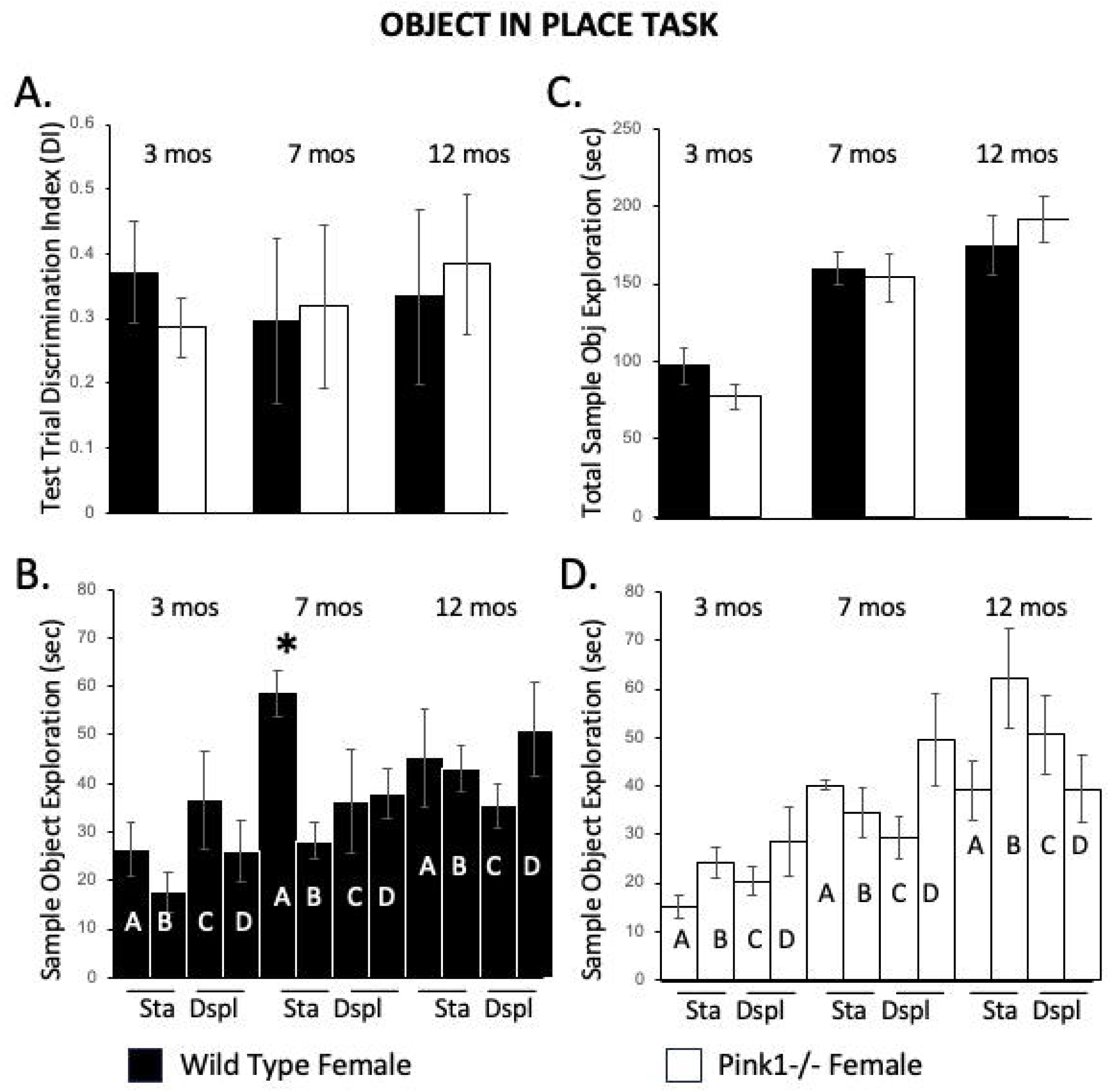
Bar graphs summarizing test trial (A) and sample trial (B-D) data for Object-in-Place testing in wild-type (black bars) and Pink1 knockout (Pink1-/-; white bars) female rats. Data from testing at 3, 5 and 7 months of age are shown. At every age tested, robust measures of discrimination index (DI) were similar in Pink1-/- and W (A). Analyses of total times spent investigating objects during sample trials also revealed no significant group differences [C, in seconds (sec)]. When sample trial exploration times were explored by individual objects (bars marked A-D) and by those that remained stationary (Sta)/in the same place or were displaced (Dspl) during the test trial (B, D), there was only one case where investigation times were uneven. This was for WT rats at 7 months of age, where animals spent significantly more time (*) with one of the objects (bar marked A) that remained stationary during the test trial (B).

Analyses of object exploration behaviors during sample trials found no significant group differences in total sample object exploration times (η^2^ = 0.006-0.13, Fig 6C) and regression analyses found no significant correlations between total sample trial object exploration and DI values measured during test trials for either group at any age (R^2^= 0.00-0.24, *p* = 0.18-0.99). Finally, within groups repeated measures ANOVAs found no evidence of significant bias among the four sample objects for Pink1-/- rats at any age (η^2^ = 0.11; 12 mos: η^2^ = 0.08, Fig 6B). However, for WT rats at 7 months of age there was a significant difference in exploration across objects [F_(3,27)_ = 3.61, p = 0.03; η^2^ = 0.29, Fig 6B]. This was driven by the WT female rats exploring one of the stationary objects significantly longer (∼40%, 20-30 sec) than the other three objects present (p = 0.002-0.30).

### 3.5 Episodic-Like (What, Where, When) Memory: NEW IN MALE AND FEMALE

#### 3.5.1 Effects of Biological Sex

Wild type male and female rats both demonstrated robust discrimination for object ‘what’ information at 3, 7 and 12 months of age (Fig 7A, 8A). In WT males, mean DI values ranged from 0.18 to 0.33, in females DIs ranged from 0.24 to 0.38 and one-sided t-tests confirmed that these DI values were all significantly to near significantly greater than zero (Table 2). Sex differences were noted in the other two memory domains. Specifically, WT males consistently showed stronger discrimination of ‘where’ (DIs 0.26-0.30) compared to ‘when’ (DIs 0.06-0.19, Fig 7A), whereas WT females showed better discrimination for ‘when’ (DIs 0.19-0.23) compared to ‘where’ (DIs −0.05-0.18, Fig 8A, see Table 2). One-sided t-tests further showed that DI values for ‘where’ were all significantly greater than zero in males but not in females and that DI values for ‘when’ were all significantly greater than zero in females but not in males (Table 2). However, repeated measures ANOVAs that included DIs for all three memory domains found no significant main effects of biological sex on episodic memory overall, and calculated effect sizes were uniformly small (3 mos: η^2^ = 0.004; 7 mos: η^2^ = 0.06; 12 mos: η^2^ = 0.02). Additional analyses of sample trial activities in rats of both sex, and those in which the data from WT females were stratified by estrous cycle are presented below along with data from sex-matched Pink1-/- groups.

**Figure 7.**
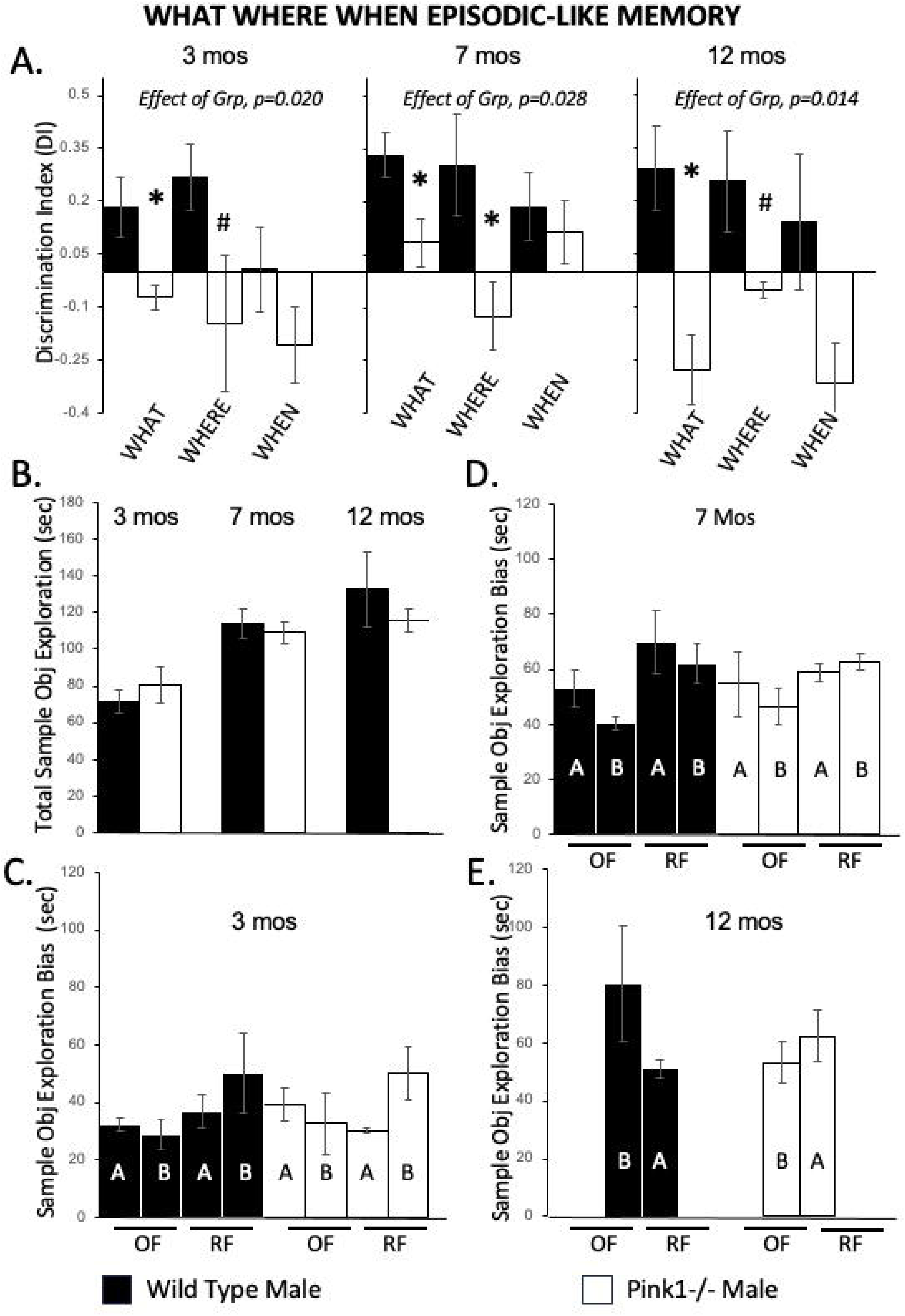
Bar graphs summarizing test trial (A) and sample trial (B-E) data for What, Where, When Episodic-Like Memory testing in wild-type (black bars) and Pink1 knockout (Pink1-/-; white bars) male rats. Data from testing at 3, 5 and 7 months of age are shown. Discrimination indices (DI) from test trials are shown in (A), along with the p value from repeated measures analysis of variance testing for main effects of group (Effect of Grp) on episodic memory across all domains. In wild type males, DI values were consistently robust for ‘what’ and ‘where’, and notably weak for ‘when’ information. In Pink1-/- males, DI values were consistently low in all three domains; DI’s for ‘what’ and ‘where’ were also near-significant (#) or significantly (*) lower than those of control males for all three testing ages. Analyses of total times spent investigating objects during sample trials also revealed no significant group differences [B in seconds (sec)]. Analyses of sample trial object exploration times where the data were stratified by trial [Sample trial 1: old familiar (OF); Sample trial 2: recent familiar (RF)] and by counterbalanced objects (bars marked A or B) used in these trials found no evidence for object bias (significant difference) within or between groups at 3 (C) or 7 months of age (D). At 12 months of age, object sets were not counterbalanced (in error); there were no significant differences within or across groups in the total amounts of time rats spent with objects during Sample trial 1 (OF) compared to Sample trial 2 (RF).

**Figure 8.**
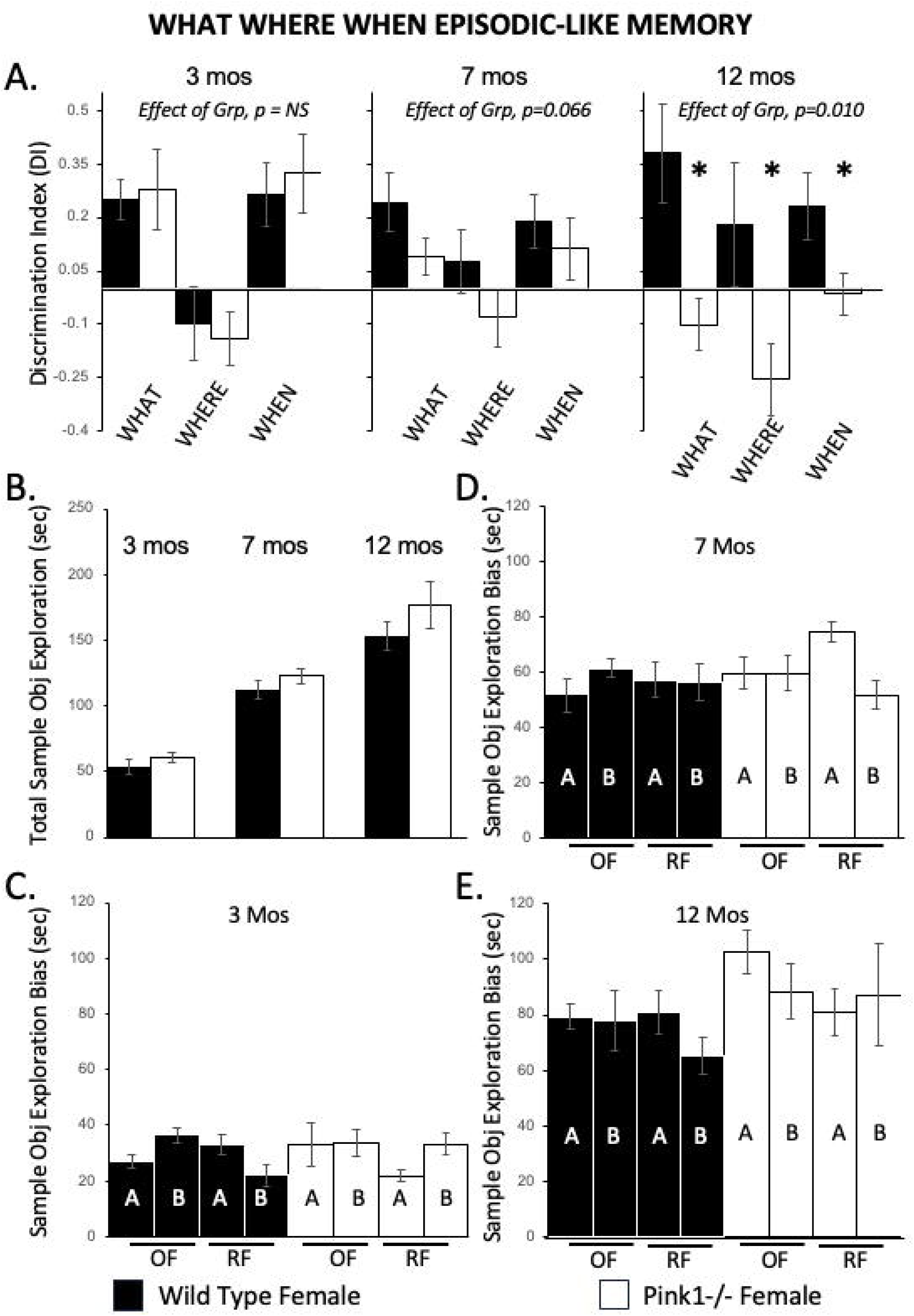
Bar graphs summarizing test trial (A) and sample trial (B-E) data for What, Where, When Episodic-Like Memory testing in wild-type (black bars) and Pink1 knockout (Pink1-/-; white bars) female rats. Data from testing at 3, 5 and 7 months of age are shown. Discrimination indices (DI) from test trials are shown in (A), along with the p-value from repeated measures analysis of variance testing for main effects of group (Effect of Grp) on episodic memory across all domains. In wild-type females, DI values were consistently robust for ‘what’ and ‘when’ and were weaker for ‘where’ information. In Pink1-/- females, DI values in all three domains were similar to wild types at 3 months of age; were lower, but not significantly lower at 7 months of age; and were significantly (*) lower than wild-type females at 12 months of age. Analyses of total times spent investigating objects during sample trials also revealed no significant group differences [B, in seconds (sec)]. Analyses of sample trial object exploration times where the data were stratified by trial [Sample trial 1: old familiar (OF); Sample trial 2: recent familiar (RF)] and by counterbalanced objects (bars marked A or B) used in these trials (C. 3 months; D. 7 months; D. 12 months) found no evidence for object bias (significant difference) within or between groups.

**Table 2.** Results from testing wild-type male and female rats on the What, Where, When Episodic-like memory task (WWW). Values of discrimination index (DI) ± s, standard error of the mean are shown, along with results from one-sided t-tests that compared group mean DI values to zero. Effect sizes (Cohen’s d, *d*) are also listed.

| **WWW: 3 Months** | **What** | **Where** | **When** |
| --- | --- | --- | --- |
| *Wild-Type Male* | DI = 0.182 ± 0.084<br>*t*(5) = 2.170, *p* = 0.041<br>*d* = 0.886 | DI = 0.267 ± 0.095<br>*t*(5) = 2.824, *p* = 0.018<br>*d* = 1.153 | DI = 0.007 ± 0.083<br>*t*(5) = 0.056, *p* = 0.479<br>*d* = 0.023 |
| *Wild-Type Female* | DI = 0.238 ± 0.056<br>*t*(9) = 4.27, *p* = 0.001<br>*d* = 1.35 | DI = -0.053 ± 0.104<br>*t*(9) = -0.507, *p* = 0.31<br>*d* = -0.16 | DI = 0.229 ± 0.090<br>*t*(9) = 2.549, *p* = 0.016<br>*d* = 0.806 |
| **WWW: 7 Months** | **What** | **Where** | **When** |
| *Wild-Type Male* | DI = 0.333 ± 0.064<br>*t*(5) = 5.224, *p* = 0.002<br>*d* = 2.133 | DI = 0.304 ± 0.144<br>*t*(5) = -2.113, *p* = 0.051<br>*d* = 0.945 | DI = 0.187 ± 0.096<br>*t*(5) = 1.953, *p* = 0.054<br>*d* = 0.797 |
| *Wild-Type Female* | DI = 0.241 ± 0.082<br>*t*(9) = 2.928, *p* = 0.008<br>*d* = 0.926 | DI = 0.152 ± 0.088<br>*t*(9) = 0.872, *p* = 0.203<br>*d* = 0.276 | DI = 0.189 ± 0.074<br>*t*(9) = 2.558, *p* = 0.015<br>*d* = 0.809 |
| **WWW: 12 Months** | **What** | **Where** | **When** |
| *Wild-Type Male* | DI = 0.292 ± 0.118<br>*t*(4) = 1.716, *p* = 0.081<br>*d* = 0.371 | DI = 0.257 ± 0.143<br>*t*(4) = 1.263, *p* = 0.138<br>*d* = 0.412 | DI = 0.059 ± 0.199<br>*t*(4) = 0.296, *p* = 0.391<br>*d* = 0.132 |
| *Wild-Type Female* | DI = 0.381 ± 0.139<br>*t*(7) = 2.734, *p* = 0.015<br>*d* = 0.967 | DI = 0.180 ± 0.175<br>*t*(7) = 1.029, *p* = 0.169<br>*d* = 0.364 | DI = 0.230 ± 0.093<br>*t*(7) = 2.469, *p* = 0.021<br>*d* = 0.873 |

#### 3.5.2 Effects of Genotype in Males

Mean DI values in Pink1-/- male rats were lower than those of WT males for all memory domains at all ages tested (Fig 7A). These observations were statistically robust. First, repeated measures ANOVAs identified significant main effects of Group (genotype) on episodic memory overall at every age [3 mos: F_(1,8)_ = 8.43, p = 0.02; η^2^ = 0.51; 7 mos: F_(1,7)_ = 7.66, p = 0.03; η^2^ = 0.52; 12 mos: F_(1,8)_ = 1.25, p = 0.01; η^2^ = 0.57]. Follow-up pairwise comparisons further showed that DI values for object ‘what’ information, which ranged in the Pink1-/- group from −0.28 to 0.09, were all significantly lower than those of WT controls [3 mos: F_(1,9)_ = 8.32, p = 0.02; η^2^ = 0.48; 7 mos: F_(1,8)_ = 6.63, p = 0.03; η^2^ = 0.45; 12 mos: F_(1,9)_ = 13.94, p = 0.005; η^2^ = 0.61]. Further, in contrast to findings in WT rats (above), one-sided t-tests showed that ‘what’ DI values in the Pink1-/- cohort were either not significantly different than zero or were significantly less than zero (3 mos: *t*(4) = −2.84, p = 0.02, *d* = −1.27; 7 mos: *d* = 0.62; 12 mos: *t*(5) = −2.80, p = 0.02, *d* = −1.140). Discrimination indices for ‘where’ also ranged from −0.15 to −0.04 in Pink1-/- males. These values were near significantly to significantly lower than corresponding DI values in WT rats [3 mos: F_(1,9)_ = 4.10, p = 0.07; η^2^ = 0.31; 7 mos: F_(1,9)_ = 6.40, p = 0.03; η^2^ = 0.42; 12 mos: F_(1,8)_ = 4.56, p = 0.07; η^2^ = 0.36] and were either not significantly greater than zero (3 mos *d* = −0.34, 7 mos *d* = −0.52) or were significantly less than zero (12 mos: *t*(4) = −2.14, p = 0.05, *d* = −0.87). Finally, although DIs for ‘when’ were low in both groups, they were lower in the Pink1 cohort (DI = −0.32 to 0.11). Group differences, however, were not significant at any age, calculated effect sizes (η^2^) ranged from 0.03-0.25, and one-sided t-tests showed that in WT and Pink1-/- males most ‘when’ DIs were not significantly different than zero (*d* = −0.86-0.79) with the exception of ‘when’ DI for Pink1-/- males at 12 mos of age, which was significantly less than zero (*t*(5) = −2.85, p = 0.02, *d* = −1.16).

Analyses of total object explorations during the two sample trials found no significant group differences in sample object exposures (η^2^ = 0.02-0.06, Fig 7B). Regression analyses also found only 2 out of 18 cases were positive correlations between sample trial object exploration and subsequent measures of DI were significant; When DI for 12-month-old WT rats (R^2^ = 0.89, *p* = 0.02); and When DI for 3-month-old Pink1-/- rats (R^2^ = 0.90, *p* = 0.01). All other correlations were non-significant (R^2^ = 0.00-0.65, *p* = 0.08-0.99). Within groups ANOVAs also found no evidence for significant bias among the counterbalanced objects used as old familiar vs. recent familiar objects (Fig 7C-E) for either group (3 mos: WT: η^2^ = 0.30, Pink1-/- η^2^ = 0.31; 7 mos: WT: η^2^ = 0.02, Pink1-/- η^2^ = 0.19; see Methods for 12 mos).

#### 3.5.3 Effects of Genotype in Females

At 3 months of age, mean DI values were similar in Pink1-/- and WT female rats for all memory domains (Fig 8A). Similar to DI values in the WT group (above) mean DI values in Pink1-/- females were robust for ‘what’ (DI = 0.28) and ‘when” (DI = 0.33) and were weak for ‘where’ (DI = −0.14). A repeated measures ANOVA confirmed that there were no significant group differences in episodic memory at this time point months [η^2^ = 0.002]. Follow-up pairwise comparisons, not technically allowed, further showed that at 3 months, there were no significant group differences for any DI values and that calculated effect sizes were small (η^2^ = 0.003-0.03). As in WT females, one-sided t-tests also identified DI values for ‘what’ and ‘when’ in Pink1-/- females as significantly greater than zero (What: *t*(9) = 2.43, p = 0.008, *d* = 0.77; When: *t*(9) = 2.94, p = 0.02, *d* = 0.93) and mean DI for ‘where’ as significantly less than zero (*t*(9) = −1.89, p = 0.05, *d* = −0.60).

At 7 and 12 months of age, episodic memories were noticeably lower in the Pink1-/- cohort (Fig 8A). Thus, mean DI values in Pink1 -/- females for ‘what’, ‘when’ and ‘where’ fell to 0.09, 0.08, and 0.11, respectively, at 7 months, and at 12 months were lower still (What: DI = −0.10; Where DI = −0.25; When: DI = −0.02). Repeated measures ANOVAs performed on these data identified group differences in episodic memory overall that were near significant at 7 months [F_(1,19)_ = 3.81, p = 0.07; η^2^ = 0.17] and significant at 12 months of age [F_(1,14)_ = 1.76, p = 0.010; η^2^ = 0.38]. Follow-up wise comparison showed that for the 7-month data, there were no significant group differences for any memory domain. However, calculated effect sizes ranged medium to large (η^2^ = 0.02-0.11) and one-sided t-tests showed that DI for ‘where’ was near significantly less than zero *t*(10) = −1.79, p = 0.05, *d* = −0.30 and DI’s for ‘what’ and ‘when’ were not significantly different than zero (What: *d* = 0.54; When: *d* = 0.39). Finally, at 12 months of age group differences in DI were all significant [What: F_(1,14)_ = 9.35, p = 0.009; η^2^ = 0.400; Where: F_(1,15)_ = 4.87, p = 0.04; η^2^ = 0.25; When: F_(1,14)_ = 4.94, p = 0.04; η^2^ = 0.26]. and one-sided t-tests showed that DIs for ‘what’ (*d* = −0.48) and ‘when’ (*d* = −0.09) were not significantly different than zero (above) and that the mean DI for ‘where’ was significantly less than zero (*t*(8) = −2.50, p = 0.02, *d* = −0.83). Finally, when the DI data from WT and Pink1-/- groups were stratified by estrous cycle stage, a significant differences was noted for DIs for What in Pink1-/- rats at 3 months at estrous stages associated with low (DI= 0.48, n = 6) compared to high circulating hormone levels (DI = −0.02, n = 4) on the day of testing [F (1,9) = 8.242, p = 0.021,η^2^ = 0.51]. However, no other significant effects of estrous cycle were found for either group for any memory domain at any age (η^2^ = 0.000-0.353). The numbers of WT and Pink1-/- subjects in estrous or proestrus vs. diestrus that were compared in each analysis are shown in Table 1.

Analyses of sample trial behaviors found no significant group differences in total sample object exploration times (η^2^ = 0.060-0.063, Fig 8B). Regression analyses did, however, find 2 instances out of 18 where there was a significant positive correlation between total sample trial object exploration time and DI [When DI for 12 month old WT rats (R^2^= 0.61, F_(1,7)_ = 9.27, *p* = 0.02) and Where DI for 7 month old Pink1-/- rats (R^2^= 0.38, F_(1,10)_ = 5.41, *p* = 0.05)]. No other correlations were significant (R^2^= 0.001-0.39, *p* = 0.05-0.93). Within groups ANOVAs also found no evidence for significant bias among counterbalanced objects used as stimuli during the first (old familiar) vs. second (recent familiar) test trials for either group at all ages (3 mos: WT: η^2^ = 0.26, Pink1-/- η^2^ = 0.15; 7 mos: WT: η^2^ = 0.11, Pink1-/- η^2^ = 0.33; 12 mos: WT: η^2^ = 0.05, Pink1-/-: η^2^ = 0.03, Fig 8C-E).

### 3.6 Olfactory Discrimination and Habituation

Differences in odor were among the features that rats could use to discriminate among sample object sets in the WWW task. Given the prevalence of anosmia/hyposmia in PD, in addition to analyses of sample object explorations (above), rats in all groups were also tested for habituation and dishabituation to non-social odors. Rats in all groups exhibited highest levels of sniffing during first encounters with each odor and markedly less sniffing during the odor’s second and third presentations (Fig 9). As expected, repeated measures ANOVAs that compared sniffing durations for each trial across sex (Male WT vs. Female WT) and within sex across genotype (WT vs. Pink1-/-) identified significant main effects of Trial (Male WT vs. Female WT: F_(8,96)_ =6.77, p < 0.001, η^2^ =0.361; Female WT vs. Female Pink1-/- F_(8,136)_ =9.53, p < 0.001, η^2^ =0.359; Male WT vs. Male Pink1-/- F_(8,72)_ =5.48, p < 0.001, η^2^ =0.378). However, there were no significant interactions between sex or genotype and Trial and no main effects of sex or genotype. Calculated effect sizes were also modest to small for factors of sex in WT rats (η^2^ = 0.192) and factors of genotype in males and females (η^2^= 0.000-0.289). A series of repeated measures ANOVAs that compared mean group sniffing times across trials where data in WT and Pink1-/- female rats were stratified by estrous cycle stage found no significant effects of estrous cycle in either group (WT: water: η^2^ = 0.02; banana: η^2^ = 0.00; almond: η^2^ = 0.18; Pink1-/-: water: η^2^ = 0.13; banana: η^2^ = 0.08; almond: η^2^ = 0.09). The numbers of WT and Pink1-/- subjects in estrous or proestrus vs. diestrus that were included in these comparisons are shown in Table 1.

**Figure 9.**
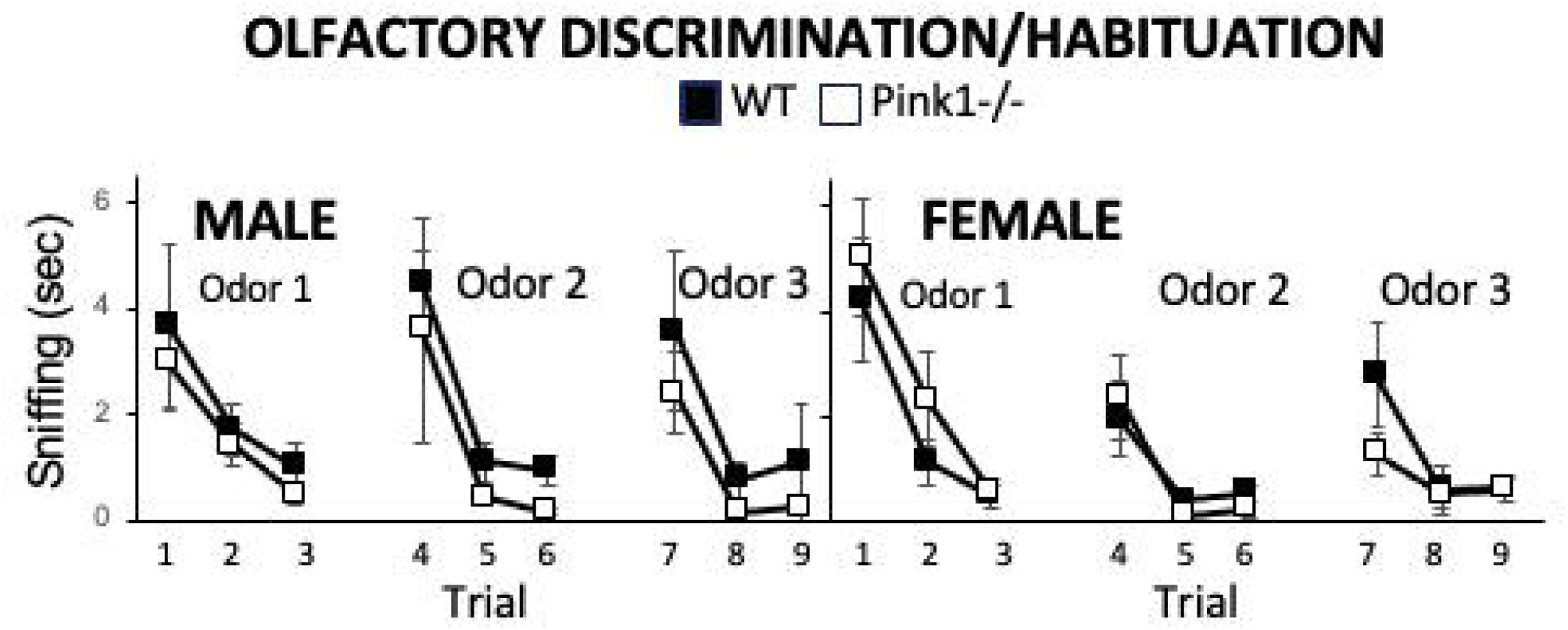
Line graphs showing the amounts of time in seconds (sec) rats spent sniffing three non-social odors that were each presented three times. Female rats (right hand graphs) spent significantly less time (*) than males sniffing Odor 2 and Odor 3 upon their first presentation. However, there were no significant effects of the Pink1 knockout (Pink1-/-, open circles) compared to wild type (black circles) on measures of odor discrimination (Trials 1, 4, 7) or habituation (Trials 1-3, 4-6, 7-9) in rats of either sex.

## 4. DISCUSSION

Mild impairments in cognition and memory manifest early to prodromally in up to 50% of patients diagnosed with PD(*87*). Many of these deficits are resistant to or worsened by the pharmacologic and non-pharmacologic interventions, e.g., deep brain stimulation, that are commonly used and highly effective in improving patients’ motor function(*7, 8, 88–91*). As a result, cognition and memory impairments in PD tend to persist, worsen over time, and exact increasingly tolls on patients, caregivers and healthcare systems. Moreover, an early presence of these non-motor symptoms signals increased risk for rapid clinical deterioration, for freezing of gait and falls and for developing PD-related dementia(*92–95*), which are all leading causes of patient disability, institutionalization and even death(*96–98*). Thus, there are urgent needs to better understand the neurobiology that underpins cognitive and memory impairments in PD and to use this information to develop better and potentially disease-modifying ways of intervening early on to treat them. For animal models to be useful in addressing these needs, they should ideally have construct validity and broad face validity for the timing and progression of PD-relevant neuropathological brain changes, non-motor symptoms and motor signs. The present studies add to the evidence showing that Pink1-/- rats meet many of these criteria.

Most studies in Pink1-/- rats have focused on pathophysiology and/or motor function. However, recent studies have shown that male Pink1-/- rats also develop significant deficits in spatial cognition/route navigation(*79*) and in several types of object-based recognition memory prior to the emergence of measurable deficits in somatic motor operations(*80*). The longitudinal behavioral testing studies reported here confirmed that recognition memories for object form (NOR) and location (NOL) are intact in Pink1-/- male rats at 3 months of age but become significantly impaired by the time rats are 5 months old. Evaluation of episodic memory (WWW)–carried out here for the first time in Pink1-/- rats, further showed that discrimination for ‘what’, ‘where’, ‘when’ domains are significantly impaired in male Pink1-/- by the time they reach 3 months of age, i.e., weeks before deficits in spatial learning and other object memory tasks are present.

Finally, behavioral testing in female Pink1-/- rats showed that these animals were unimpaired in NOR, NOL or OIP in testing through 12 months of age. However, in WWW testing, DI values were only similar to WT females at 3 months of age. By 7 months of age, DI values for all domains were noticeably but not significantly worse than controls age and by 12 months of age WWW performance in female Pink1-/- rats was significantly worse than WT females in all three episodic memory domains. As discussed below, these data bring to light new ways in which Pink1-/- rats model the so-called mild cognitive deficits associated with PD, including the major sex differences that characterize them as well as the more nuanced clinical features that distinguish deficits in episodic memory from other at-risk cognitive or memory domains.

### 4.1 Cognitive Deficits in Pink1-/- Rat Models of PD: Previous Studies and Findings from Object Recognition Memory Tasks

Animal models play critical roles in identifying the neurobiological causes and consequences of PD and in developing safer, more effective ways to treat them. As reviewed in the Introduction, the early losses of cognitive and memory function in PD have been most frequently investigated in rodent models where disease-relevant pathophysiology is induced by selective chemical lesions, exposures to environmental toxins or by intracerebral injections of abnormally folded protein fibrils where dosages and/or delivery strategies have been designed to emulate pre-motor stages of disease(*40–42, 99*). While findings to date largely affirm that these models recapitulate cognitive and memory deficits as well as some of their differences in male and female PD patients, their acute or semi-acute timelines have limited capacity to address questions related to the arcs of disease. These include the early to prodromal onsets and progressions of non-motor symptoms and their functional and temporal relationships to later developing motor signs. Being able to investigate these dynamic aspects of PD could lead to discoveries that enable earlier diagnosis of disease and the development and testing of treatments that may be preventive, disease-altering and/or unfettered by potentially negative consequences of treating motor signs for non-motor symptoms, and vice versa. Rodent models that have been genetically engineered to carry PD-relevant risk alleles or gene mutations are well-suited for these sorts of studies. As discussed below, among the available models, Pink1-/- rats appear to hold significant benefits for analyses that are focused on non-motor symptoms involving cognition and memory.

Several findings link the loss of Pink1 and PINK1 function to cognitive impairment. For example, Pink1 function has been shown to be diminished in the cerebral cortex and hippocampus of mouse models of Alzheimer’s disease(*100–102*), and restoration of Pink1 function in these models has been shown to reduce amyloid deposition and improve spatial memory in Morris Water Maze performance(*101, 103*). Patients in which PD is causally linked to loss of function *Pink1* mutations have also been shown to be the most likely of all monogenetic forms of disease to develop deficits in cognition and memory(*67, 104*). Since their early characterization(*66*), studies in Pink1-/- rats have mainly focused on cellular and molecular pathologies and on progressive deficits in somatic and otolaryngeal motor function. However, one of the earliest studies to behaviorally screen this strain for deficits in cognition or memory concluded that there were no deficits in NOR or Barnes maze performance in 6-8 month old Pink1-/- males(*105*). It is important to note, however, that likely due to the use of a single sample trial in NOR testing (see below), the WT group spent equal or less time exploring the novel compared to familiar test trial objects(*105*). Further, although Barnes maze testing was described as spanning multiple daily trials and multiple testing days, the single heat maps shown (WT and Pink1-/-) did not identify which trial(s) were represented in the images and provided no quantitative or other data to support negative conclusions(*105*). These factors could explain the contrast between the lack of cognitive phenotype reported in these studies and findings of significant deficits in object recognition memory and spatial learning that are more recently reported for male Pink1-/- rats.

Behavioral testing in male Pink1-/- rats in this lab and others has shown that at 3 (*80*) or 4 months of age(*79*), DI values for NOL, OIP and /or NOR were all robust in knockout rats and similar to measures of discrimination observed in the WT rats. However, by 5 months of age, performance in all three object recognition memory tasks dropped to and remained at levels of discrimination that were significantly lower than sex-matched controls and that were not significantly different from zero, i.e., indicating no preference for unfamiliar, moved or rearranged test objects (*80*). The present studies re-examined NOR and NOL in new cohorts of Pink1-/- and WT male rats at 3 and 5 months of age and replicated these key findings. Specifically, performance in both tasks was indistinguishable from WT in Pink1-/- males at 3 months of age and was significantly impaired two months later (5 months of age). Studies examining spatial learning have also shown that 4-month-old Pink1-/- male rats have significant difficulties in recalling previously learned complex maze routes(*79*). In comparison to WT rats, the Pink1-/- male rats took significantly longer to traverse the series of T-junctions that made up the maze and made more errors, i.e., entered incorrect arms at junction points, along the way(*79*). Together, these data demonstrate that male Pink1-/- rats spontaneously develop deficits in aspects of cognition and memory that are similar to those that are most at risk in male patients diagnosed with PD. Rats are considered adults or young adults at 4-5 months of age. Thus, the age of onset is congruent with the early onset of PD symptoms in patients carrying loss of PINK1 mutations(*106*). As in patient populations as well, these cognitive and memory symptoms manifested prior to the onset of limb weakness, uncoordinated gait and other measures of somatic motor impairment(*66, 80*). A further contribution of the present study was to identify face validity in the Pink1-/- rat model for the sex differences that distinguish the so-called mild cognitive impairments of PD.

This study is the first to rigorously examine cognition and memory in female Pink1-/- rats. Longitudinal behavioral testing in these animals showed that female Pink1-/- rats are protected from the early appearing, enduring object recognition memory deficits that impact Pink1-/- males. Specifically, testing in female Pink1-/- rats from 3 through 12 months of age showed that DI values for NOR, NOL and OIP were consistently robust, significantly greater than zero and similar to corresponding performance measures of WT females. The only case where a significant difference was noted between DI values of WT vs. Pink1-/- females was for NOR testing at 3 months of age; in this instance, DI values were notably strong in both groups, but were exceptionally high in WT females (WT =0.52; Pink1 = 0.31). These data are consistent with the behavioral sparing observed for object-based recognition memory testing in female rats where PD is modeled by 6-OHDA dopamine lesions and monoamine depletions (*50, 52*). However, the present results show that these relative protections persist through 12 months of age. This is well past the ages when motor and other anomalies develop in female Pink1-/- rats(*76, 107–109*), and past the age when endocrine and neuroendocrine changes associated with rodent reproductive senescence are expected(*110, 111*).

Although under-powered, analyses where the data were stratified by estrous cycle stage also yielded no clear evidence for performance in Pink1-/- females varying as a function of higher or lower levels of circulating ovarian hormone levels on testing days. Although further studies are needed, provisional conclusions are that protections of object recognition memory functions in female Pink1-/- rats may not be ovarian hormone dependent. Rather than contradict clinical evidence for increased vulnerabilities of cognition and memory in female PD patients as they age(*38, 112–114*), these findings appear to align with earlier studies in PD rat models suggesting that impacts of sex hormones on object-based recognition memory functions are complex, task-specific and, at least in males, are in some cases hormone-independent(*50*).

Although these assertions need to be independently established for female rats, the findings presented below for late-emerging deficits in episodic memory in female Pink1-/- rats may provide further support for the premise of task specificity in the impacts of gonadal steroid hormones on cognitive vulnerability in PD and in rat models of PD. Further discussion of these and other possibilities are preceded, however, by a brief consideration of the protocol modifications that were needed to optimize episodic memory testing in Long Evans rats.

### 4.2 Modifications to the WWW Paradigm

Previous studies have shown that Long Evans rats can be susceptible to neophobia and novel object avoidance (*115, 116*). This presents challenges to object-based recognition memory testing-where outcome measures are based on positive expressions of preference for novelty, e.g., object approach and exploration (*117, 118*). However, previous studies have shown that the tendency for novel object avoidance in this rat strain are mitigated specifically by using three sequential sample trials rather than single exposures to sample objects(*115, 116*). This is easily adaptable to NOR, NOL and OIP testing and implementation of these protocols in the present and previous studies(*80*) have yielded reliable, reproducible results in WT and Pink1-/- rats of both sexes. However, standard versions of the WWW task(*119, 120*) require single, sequential presentations of two distinct sample object arrays prior to a third and final test trial. This trial structure is not amenable to the use of repeated sample exposures. As expected, pilot studies conducted in this lab in WT Long Evans males and females using the standard Dere version of the WWW task produced variable sample trial object exploration behaviors and inconsistent test trial data that did not reliably reflect episodic memory status (data not shown). In efforts to encourage rats’ approach and exploration of all four sample and all four test trial objects presented, distinct odor cues were added to the tactile, visual and other features that distinguished the quadruplicate object sets. Pilot studies were conducted in which odorants were placed inside of objects that had small holes in the top surface, e.g, saltshakers. These confirmed that rats could detect but not lick, taste or eat the odorant; that WT male and female Long Evans rats approached scented objects more quickly than unscented objects; that rats tended to explore all four objects present; and that rats not only sniffed the open ports, but also actively explored other features of the sample objects. These object modifications also resulted in consistent expressions of expected patterns of test trial object discriminations. Thus, they were adopted here to explore ‘what’, ‘where’ and ‘when’ episodic memory domains in male and female Pink1-/- rats.

In carrying out these studies, several cautions were kept in mind. First, although odor was not the only dimension that rats could use to distinguish object sets, it did serve as an enticement. Thus, it was essential to ensure that the odorants themselves did not introduce unintended non-mnemonic biases in exploratory behavior. This was done in several ways. First, pairs of odorants that fell into the same class (spice, sweet or savory) were used for old familiar (Sample Trial 1) and recent familiar (Sample Trial 2) objects. Further, the object/odor combinations used were counterbalanced across subjects within all groups. To detect potentially confounding effects of odor bias, total sample trial exploration times and test trial DI values for what, where and when domains were quantitatively compared for rats where one or other object/odor combinations were used both within and across groups. None of these assessments revealed instances where odor biases were likely to have significantly swayed–either positively or negatively, test trial performance data. A further consideration, however, is that anosmia/hyposmia occurs in an estimated 45-90% of patients with PD and that these deficits are among the earliest manifestations of disease of familial and sporadic forms of PD(*121–123*). Although olfactory deficits have been reported in other rodent models of PD, previous studies of sensitivity to new odors in 3-month-old Pink1 -/- rats showed no such deficits(*124*). In the present studies, objective tests of olfactory discrimination of and habituation to non-social odors performed in all groups at 5-6 months also showed no significance differences among WT and Pink1-/- rats of either sex in detecting, differentiating and adapting to the sequential presentations of three non-social odorant stimuli (water, almond extract diluted in water and banana extract diluted in water).

Thus, the addition of distinguishing odors to WWW sample objects yielded intended effects of encouraging object exploration without introducing unintended, non-mnemonic effects on episodic memory measures. It may nonetheless be advisable for future studies to test whether adding the same odor to visually or otherwise distinguishable sample object sets also mitigates neophobia, as this would avoid the potential for different olfactory cues to be of mis- or un-matched valence.

### 4.3 Cognitive Deficits in Pink1-/- Rat Models of PD: Episodic Memory for ‘What’, ‘When’ and ‘Where’

The first higher order behavioral constructs to become impaired in PD typically involve memory function and episodic memory function in particular (*9, 125–130*). These deficits in recollection of contextualized life events are also a common cognitive complaint at the time of clinical diagnosis(*10, 81, 82, 131, 132*), and their presence in early stages of disease can be uniquely predictive of faster rates of clinical decline, greater symptom severities and greater likelihood of developing PD-related dementia(*92–94, 133, 134*). Similar to other cognitive and memory domains, sex differences have been identified in the prevalence and treatment response of certain types of episodic memory losses in PD. Male patients, for example, have been shown to be more often and/or more severely impaired in episodic memory for verbal and visuospatial information (*26, 31*). Given their early/earliest compromise, understanding the neural bases for episodic memory deficits–and their sex differences, could be valuable in facilitating earlier diagnosis of illness and developing targeted treatments that are preventive or disease-modifying. Accordingly, episodic memory functions have been and continue to be subjects of intensive clinical study. These include behavioral studies parsing the impacts of PD on discrete components of episodic memory, e.g., information encoding, storage and retrieval(*135–137*), and imaging studies examining network and other brain correlates, e.g., cortical thinning, hippocampal atrophy, white matter abnormalities, of episodic memory impairment(*138–142*). However, these studies mainly involve symptomatic PD patients. Validated animal models, on the other hand, could be useful in pushing windows of investigation back to earlier and ideally prodromal stages of disease. This in turn could accelerate the discovery of biomarkers and/or therapeutic strategies that target at-risk cells and circuits before they become significantly and perhaps irreversibly compromised by the progression of disease. To our knowledge, however, there has been only one formal investigation of episodic-like memory in a rodent model of PD.

Episodic memory has been previously investigated in adult male and female rats in which premotor stages of PD were modeled by partial neostriatal 6-OHDA dopamine lesions(*143*). Using the Dere version of the What, Where When Episodic-like Memory task(*120*), this single time point study showed that 6-OHDA lesions profoundly impaired all three episodic memory domains in both male and female rats(*143*). It is important to recall, however, that the same and similar PD models also produce deficits in males but not females in other object recognition-based memory tasks(*50*). This suggests that 6-OHDA lesions may affect the neural circuits that underlie episodic differently from those associated with, for example, performance in NOR and OIP tasks. However, the compressed time frame over which lesion-induced neural changes occur makes it difficult to parse impacts across time and thus, across brain circuits. The present findings suggest that Pink1-/- rats on the other hand can be important in making these differentiations.

As discussed above, object recognition memories (NOR, NOL, OIP) are intact in male Pink1-/- rats when animals are 3-4 months of age(*79, 80*). However, performance in all three tasks are significantly impaired weeks later when rats are about 5 months old. The present studies now show that significant deficits in ‘what’, ‘where’ and ‘when’ domains of episodic memory manifest 1-2 months earlier than these other indices of memory impairment and are already present in Pink1-/- males at 3 months of age. As for other memory tasks, once established, significant deficits in WWW persist and are present in Pink1-/- males through testing at 12 months of age. A very different pattern of vulnerability in episodic memory was observed in female Pink1-/- rats. Thus, at 3 months of age, Pink1-/- female rats showed no discernable deficits in any domain of episodic memory. However, at 7 months old, WWW DI values in the knockout group were visibly but not significantly lower than those of WT females. By 12 months of age, WWW performance in Pink1-/- females had declined further, yielding DI values for all three episodic memory domains that were significantly lower than WT controls and that were not significantly different than zero. These sex-specific findings in Pink1-/- rats align with clinical data that identify episodic memories as uniquely vulnerable in PD. First, in keeping with clinical evidence for the early vulnerability of episodic in PD(*81, 82, 131*), it is notable that WWW deficits were detected in Pink1-/- rats of both sex prior to the onset of all other cognitive and memory impairments described in this strain to date(*79, 80*). In addition, in Pink1-/- males, prodromal WWW deficits were immediately profound and enduring. This along with the later-but still prodromal, emergence of deficits in other classes of cognitive and memory function suggests that male Pink1-/- rats could be useful in identifying the neural substrates that underpin the predictive power of early appearing episodic memory deficits for rapid clinical decline and risk of developing PD-related dementia(*93, 133*). The later occurring disturbances in episodic memory seen in female Pink1-/- rats are also in keeping with clinical evidence for earlier vulnerability of episodic memories and other behavioral constructs in males diagnosed with PD(*26, 31*) and for increasing the susceptibility of female PD patients to deficits in cognition and memory as a function of age(*38, 112–114*). Because the later-life vulnerability of female patients to higher-order deficits in PD coincides with reproductive senescence, these clinical observations have been cornerstones for hypotheses that causally link declining ovarian hormone levels of cognitive or memory loss and that predict therapeutic benefits of hormone replacement(*35, 38, 144*). However, these hypotheses have been difficult to prove in clinical studies alone. The timelines of onset suggest that studies of WWW deficits in female Pink1-/- rats could be important in clarifying the impacts of age-related changes in endocrine and neuroendocrine status on cognition and memory in female PD patients and optimizing hormone-based interventions. One strategy could be to incorporate the collection of tail vein blood into the behavioral testing regimen to probe for quantitative relationships between episodic memory function and multiple measures of hormone and health status. Finally, while the merits of Pink1-/- rats as models of the PD prodrome have been well articulated(*74*), the present findings in females suggest that the utility of this genetic rat strain may also extend to menopause, andropause and other milestones of human aging when cognition and memory are especially and differentially vulnerable in male and female PD patients.

## 5. Conclusions

An estimated 40-90% of patients with PD will experience deficits in cognition and memory(*8, 9, 126*). In most cases these impairments appear early in the course of illness and are refractory to or worsened by available treatments(*7, 88, 90, 91, 145*). Thus, they represent enduring, chronically disabling aspects of PD that significantly detract from patients’ activities and quality of life(*146–148*). This brings real urgency to the need to identify the neural bases of these symptoms and develop better ways to treat them. Findings of greater susceptibilities among male patients to cognitive deficits[(*19, 31, 149*) and of sex differences in treatment response (*150, 151*)are just some of the evidence pointing to biological sex as being key components to both endeavors. While related hypotheses for hormone protections and for benefits of hormone augmentation therapies continue to be investigated clinically(*38, 39, 152*) the data presented here identify Pink1-/- rats as an appropriate translational research tool to accelerate the pace of discovery. First, these studies confirmed that in male Pink1-/- male rats object recognition memory processes tapped in NOR, NOL are intact at 3 months of age but are impaired by the time rats are 5 months old. They further showed that enduring deficits in episodic memory emerge in male Pink1-/- rats up to 2 months earlier than all other signs of cognitive or memory impairment identified in this strain to date. Finally, the first longitudinal evaluation of cognition and memory in female Pink1-/- rats showed a lasting resilience to memory constructs measured in NOR, NOL and OIP tasks and late emerging, slowly progressing deficits in episodic memory that fully manifested in rats at around 12 months of age. Although no animal model represents all key features of PD, these data identify important strengths of Pink1-/- rats for broadly modeling sex differences in the prevalence, timing and severity of cognitive and memory deficits in PD, and for recapitulating some of the key clinical features that set vulnerability of episodic memories apart from other cognitive or memory processes that are at risk in PD. Future studies in which the benefits of this strain may be leveraged could include the following. First, the significant time lag between the onset of episodic vs. other cognitive and memory deficits in Pink1-/- males could allow activity mapping, chemogenic or optogenetic stimulation/inhibition or other strategies to be used to pinpoint the earliest circuits and cellular mechanisms to be functionally compromised in PD. Because both motor and non-motor symptoms are expressed, this strain could also be used to develop and test therapeutic strategies—including those involving hormone augmentation, that avoid common conundrums of treatments for motor signs being net negatives for non-motor symptoms or vice versa. Importantly, the data here also demonstrate that Pink1-/- rats provide sex specific contexts for these and other studies, which are essential frameworks for understanding the nature of PD and for optimizing care and clinical outcomes for male and female patients.

## 6. Declaration of Generative AI and AI-assisted technologies in the writing process

No generative AI and AI-assisted technologies were used in the writing process.

## 7. Acknowledgements

The authors thank Ms. Claudia Pinizzotto for assistance in animal testing and animal handling at the beginning of this study and for some assistance with data.

## 8. CRediT Authorship Contributions

**Aveena M. Desai**: Data curation, Formal analysis, Investigation, Visualization, Writing-review& editing

**Oluwagbohunmi A. Aje**: Data curation, Formal analysis, Investigation, Visualization, Writing-review& editing

**Mary F. Kritzer**: Conceptualization, Formal analysis, Funding acquisition, Investigation, Project administration, Writing-original draft, review& editing

### Funding

This work was supported by a Pilot Award from the Thomas Hartman Parkinson’s Research Center (to MFK).

